# Loss of Ubiquitin Ligase STUB1 Amplifies IFNγ-R1/JAK1 Signaling and Sensitizes Tumors to IFNγ

**DOI:** 10.1101/2020.07.07.191650

**Authors:** Georgi Apriamashvili, David W. Vredevoogd, Oscar Krijgsman, Onno B. Bleijerveld, Maarten A. Ligtenberg, Beaunelle de Bruijn, Julia Boshuizen, Daniela D’Empaire Altimari, Nils L. Visser, James D. Londino, Maarten Altelaar, Daniel S. Peeper

## Abstract

Despite the success of immune checkpoint blockade (ICB) most patients fail to respond durably, in part owing to reduced interferon gamma (IFNγ) sensitivity. Thus, elevating tumor IFNγ-receptor 1 (IFNγ-R1) expression to enhance IFNγ-mediated cytotoxicity is of potential clinical interest. Here, we show that increased IFNγ-R1 expression sensitizes tumors to IFNγ-mediated killing. To unveil the largely undefined mechanism governing IFNγ-R1 expression, we performed a genome-wide CRISPR/Cas9 screen for suppressors of its cell surface abundance. We uncovered STUB1 as key mediator of proteasomal degradation of the IFNγ-R1/JAK1 complex. STUB1 inactivation amplified IFNγ signaling, thereby sensitizing to cytotoxic T cells, but also inducing PD-L1. STUB1 loss in a rational combination with PD-1 blockade strongly inhibited melanomas *in vivo*. Clinically corroborating these results, a *STUB1*-KO gene signature was strongly associated with anti-PD-1 response. These results uncover STUB1 as pivotal regulator of IFNγ tumor signaling and provide a rationale for its inhibition combined with anti-PD-1.

## Introduction

Although immune checkpoint blockade (ICB) has been a major clinical success in the treatment of a variety of cancer indications, the majority of patients fail to show durable clinical responses^1,2^. This is caused by both upfront and acquired resistance mechanisms^3–7^, for which predictive biomarkers are being actively sought^8–17^. A common resistance mechanism relates to the insensitivity that tumors develop against cytokines secreted by cytotoxic T cells, including IFNγ and TNF^4,5,18,19^. IFNγ can promote antitumor activity indirectly, by inducing secretion of lymphocyte-attracting chemokines such as CXCL9, CXCL10 and CXCL11 and by skewing the attracted immune infiltrate to be more inflammatory. Conversely, IFNγ can inhibit tumorigenesis directly, by improving antigen processing and presentation, and by inducing the expression of cell cycle inhibitors, such as p21^Cip1^, and pro-apoptotic proteins, such as caspase 1 and caspase 8^20,21^. Moreover, IFNγ can sensitize tumor cells to other T cell-derived effector cytokines by increasing the expression of FAS and TRAIL receptors^22,23^.

In line with these biological functions, expression of IFNγ response genes in tumors is associated with better responses to immunotherapy^17,24^. These clinical findings are underscored by preclinical research showing a critical role for IFNγ in hindering tumorigenesis and maintaining tumor control^25^. Conversely, aberrations in the IFNγ response pathway, such as inactivation of JAK1, are associated with resistance to immunotherapy^4,5,18^.

Although the IFNγ signaling pathway has been studied extensively, and different regulatory mechanisms of this pathway have been uncovered, less is known about the cell-autonomous regulation of the IFNγ receptor 1 (IFNγ-R1), the essential ligand-binding receptor chain for IFNγ. Multiple experimental and clinical approaches have identified that tumor cells benefit from either loss or reduction in IFNγ-R1 levels in the context of ICB therapy^5^ or T cell pressure^6,26,27^.

However, to our knowledge the converse has not been studied. Specifically, the possibility that tumor cells with high (or induced) IFNγ-R1 expression show increased sensitivity to IFNγ-induced cytotoxicity has remained untested. Whereas disruption of IFNγ signaling is an established cancer trait contributing to immune escape, a scenario in which increased IFNγ signaling would lead to increased T cell sensitivity may be of clinical interest. This is the first question we addressed in this study. The answer prompted a second one, namely, which mechanisms govern the expression of IFNγ-R1. To address this, we performed a genome-wide CRISPR/Cas9 knockout screen. Lastly, we translated our findings to a preclinical setting, demonstrating their therapeutic and clinical relevance.

## Results

### High IFNγ-R1 expression results in increased sensitivity of tumor cells to T cell killing

Whereas it is established that loss of the IFNγ-R1 ablates IFNγ tumor signaling^5,25^, it is unknown whether the converse is also true. To assess whether increased abundance of IFNγ-R1 augments the susceptibility of tumor cells to cytotoxic T cells, we took advantage of the heterogeneity we observed for its expression levels in the human melanoma cell line D10. We FACsorted tumor cells with high and low expression levels of IFNγ-R1 (**Fig. 1a and b**). As a control protein, we determined the expression of another cell surface protein, PD-L1, which was expressed identically in the IFNγ-R1^High^ and IFNγ-R1^Low^ cell populations (**Fig. 1c**). We then investigated whether IFNγ-R1^High^ and IFNγ-R1^Low^ cells differentially responded to IFNγ. By flow cytometry, we observed that IFNγ-R1^High^ cells induced PD-L1 to a greater extent upon IFNγ treatment than IFNγ-R1^Low^ cells. This result indicates that the expression levels of the endogenous IFNγ-R1 protein dictate the strength of the response to IFNγ (**Fig. 1c**). This effect had also a biological consequence: in a competition experiment, IFNγ treatment was two-fold more toxic to IFNγ-R1^High^ than to IFNγ-R1^Low^ cells (**Supplementary Fig. 1a and b**).

**Figure 1:**
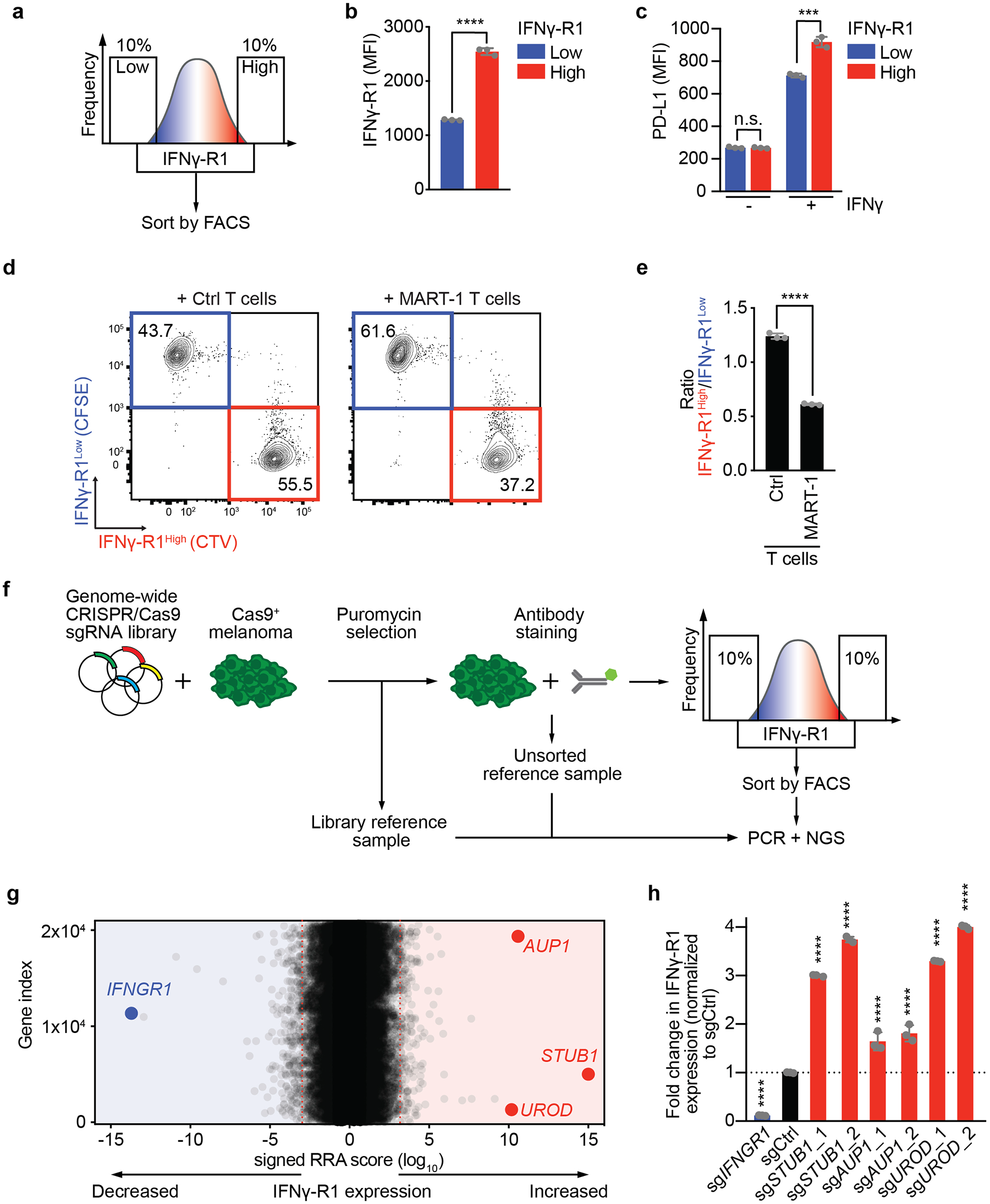
Genome-wide CRISPR/Cas9 knockout screen identifies negative regulators of IFNγ-R1 expression to modulate its cell surface abundance. **a**, Schematic outline of the FACsorting strategy to establish IFNγ-R1^High^ and IFNγ-R1^Low^ D10 human melanoma cell populations. **b**, Mean Fluorescence Intensity (MFI) of IFNγ-R1 expression on D10 melanoma cells two days after sorting the cells by flow cytometry into IFNγ-R1^High^ and IFNγ-R1^Low^ subpopulations. **c**, Assessment of IFNγ-induced PD-L1 expression of IFNγ-R1^High^ and IFNγ-R1^Low^-sorted cell populations 24 hours after treatment with 10 ng/ml IFNγ. **d**, Flow cytometry plot of the *in vitro* competition assay of IFNγ-R1^High^ vs. IFNγ-R1^Low^ cells co-cultured with either MART-1 or Ctrl T cells. **e**, Quantification of the ratio IFNγ-R1^High^ : IFNγ-R1^Low^ in competition assay of (**d**). **f**, Schematic outline of the FACsort-based genome-wide CRISPR-KO screen to identify genes regulating IFNγ-R1 cell surface expression. **g**, Screen results; red dotted lines indicate FDR cutoff <0.25 for genes enriched in 10% of cells with the highest (right) or lowest (left) IFNγ-R1 expression, as calculated by MAGeCK analysis. Gene names indicate top enriched sgRNAs in cells with the 10% highest IFNγ-R1 expression (right), as well as the sgRNAs targeting *IFNGR1* (left), serving as a positive control. **h**, Quantification of IFNγ-R1 expression by flow cytometry on cells expressing the indicated sgRNAs, plotted as fold-change in IFNγ-R1-MFI relative to sgCtrl-expressing cells. Mean±SD in (**b**), ****p<0.0001, unpaired t-test for three biological replicates. Mean±SD in (**c**), ***p=0.000467, n.s. p=0.806896, unpaired t-test for three biological replicates. Mean±SD in (**e**): ****p<0.0001, unpaired t-test for three biological replicates. Mean±SD in (**h**): ****p<0.0001, ordinary one-way ANOVA for three biological replicates with Dunnett post hoc testing.

We repeated this experiment with cytotoxic T cells, employing the matched tumor HLA-A*02:01^+^/MART1^+^ and 1D3 TCR T cell system we previously developed^19^. In this experiment also, IFNγ-R1^High^ melanoma cells showed higher susceptibility to T cell killing than IFNγ-R1^Low^ cells (**Fig. 1d, e**). Thus, the expression level of IFNγ-R1 is heterogeneous even in an established tumor cell line. More importantly, these results demonstrate that this variation has a biological consequence, in that higher IFNγ-R1 expression results in increased sensitivity of tumor cells to T cell killing.

### Whole genome CRISPR/Cas9 screen identifies regulators of IFNγ-R1 expression

Because this observation could have potential therapeutic relevance, it was important to first dissect the mechanism governing IFNγ-R1 expression in an unbiased fashion. To identify novel regulators of cell surface-expressed IFNγ-R1, we performed a CRISPR/Cas9 knockout screen (**Fig. 1f**). Cas9-expressing human D10 melanoma cells were lentivirally transduced with a genome-wide knockout library^28^, in duplicate. After two days of puromycin selection, we harvested a library reference sample. After an additional eight days of culturing, we FACsorted both the lower (IFNγ-R1^Low^) and upper (IFNγ-R1^High^) 10% of IFNγ-R1-expressing cell populations (as well as an unsorted bulk reference sample, **Fig. 1f**). Genomic DNA was isolated and sgRNA sequences were amplified by PCR. Analysis of the DNA sequencing data revealed a strong correlation between biological replicates (**Supplementary Fig. 1c**). By comparing the library reference with unsorted control samples, we confirmed significant depletion of known essential genes^29^ (**Supplementary Fig. 1d**). These quality control measures illustrate the robustness of the screen.

By MAGeCK analysis^30^, we identified several hits affecting IFNγ-R1 expression (**Fig. 1g**). Comparative analysis of the IFNγ-R1^High^ and IFNγ-R1^Low^ melanoma populations revealed that cells carrying sgRNAs targeting *IFNGR1* were most abundant in the latter population, again demonstrating the robustness of the screen (**Fig. 1g**). More interestingly, the E3 ubiquitin ligase STIP1 homology and U-box containing protein 1 (STUB1, also known as CHIP and encoded by *STUB1*) was found as the strongest hit suppressing IFNγ-R1 cell surface abundance. We also identified other genes negatively affecting IFNγ-R1 expression, including Ancient ubiquitous protein 1 and Uroporphyrinogen Decarboxylase (encoded by *AUP1* and *UROD*, respectively). We performed the same IFNγ-R1 regulator screen in a second human melanoma cell line, SK-MEL-23, which was similar in quality (**Supplementary Fig. 1e**) and also identified STUB1 and UROD (**Supplementary Fig. 1f**).

To validate these screen hits, we inactivated either *STUB1, AUP1* or *UROD* using two independent sgRNAs for each gene. Whereas cells expressing sg*IFNGR1* showed a near-complete loss of IFNγ-R1 expression, inactivation of either *STUB1* or *UROD*, and to a lesser extent *AUP1*, instead resulted in a robust increase of IFNγ-R1 abundance (**Fig. 1h**).

### STUB1 specifically regulates the cell surface fraction of IFNγ-R1

To determine whether STUB1 functions as a negative regulator of IFNγ-R1 expression beyond melanoma, we depleted it by Cas9-mediated knockout from cell lines originating from different tumor indications, and assessed the effect on the expression of IFNγ-R1. We again observed strong induction of IFNγ-R1 expression in all cell lines tested, indicating that STUB1 has a key role in limiting IFNγ-R1 expression across different tumor types (**Fig. 2a and b**).

**Figure 2:**
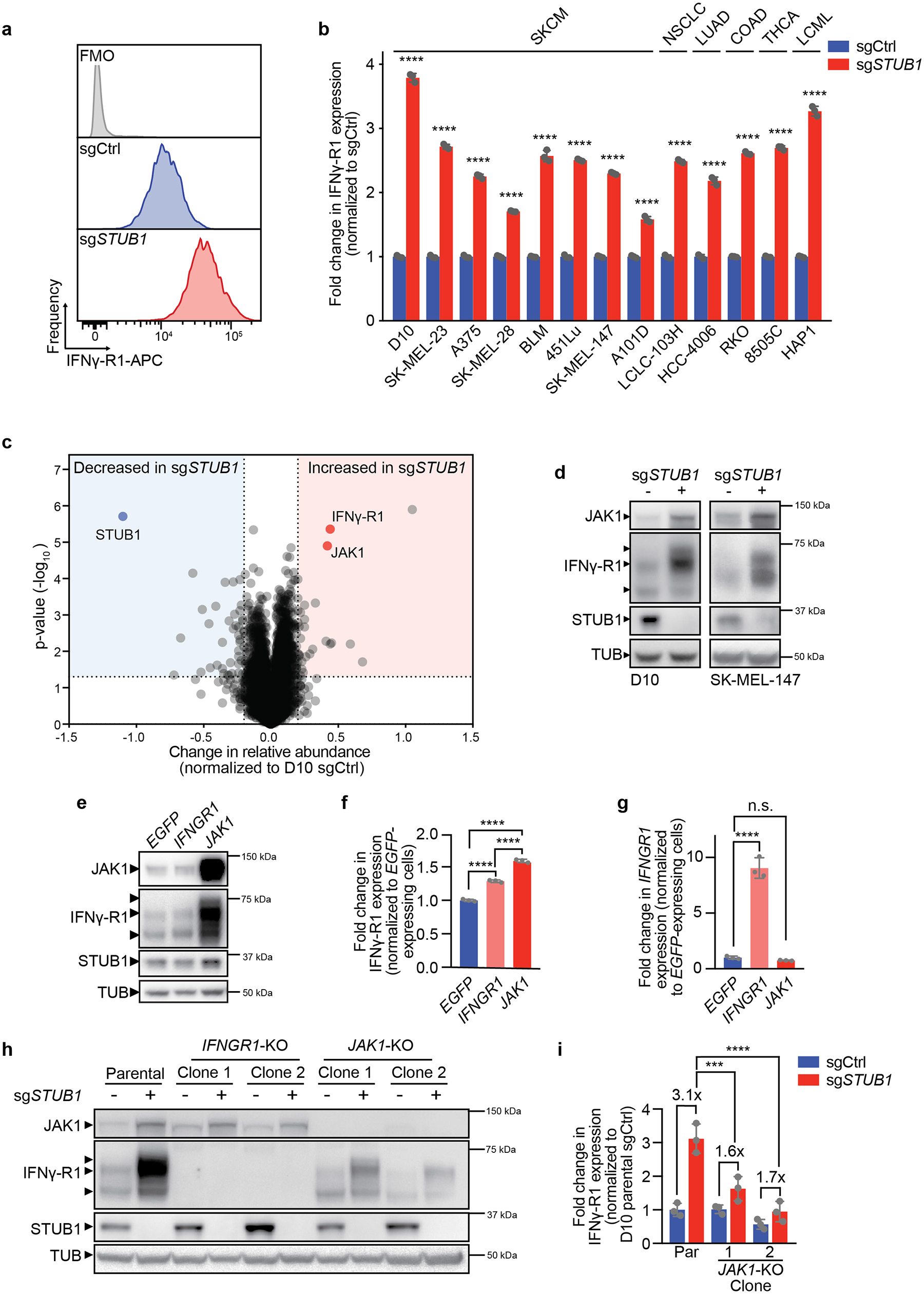
STUB1 destabilizes cell surface IFNγ-R1 in JAK1-dependent and JAK1-independent manners. **a**, Histograms of IFNγ-R1 expression on D10 melanoma cells as measured by flow cytometry in cells expressing the indicated sgRNAs. FMO: fluorescence minus one, APC: Allophycocyanin. **b**, Relative IFNγ-R1 expression (normalized to each respective sgCtrl) measured by flow cytometry in indicated human tumor cell lines expressing either sgCtrl or sg*STUB1*. Cancer types of the cell lines are abbreviated as follows: SKCM, skin cutaneous melanoma; NSCLC, non-small-cell lung cancer; LUAD, lung adenocarcinoma; COAD, colon adenocarcinoma; THCA, thyroid carcinoma. **c**, Results of proteomic profiling of D10 melanoma cells expressing either sgCtrl or sg*STUB1*. Highlighted are the top differentially regulated proteins shared between sgCtrl and sg*STUB1*-expressing D10 and SK-MEL-147 cells (**Supplementary Fig. 4e**). **d**, Immunoblot of D10 (left) and SK-MEL-147 (right) melanoma cells lines expressing either sgCtrl or sg*STUB1*. Whole cell lysates were immunoblotted for the indicated proteins (TUB is tubulin). **e**, Immunoblot of D10 melanoma cells ectopically expressing either *EGFP* (control), *IFNGR1* or *JAK1*. Whole cell lysates were immunoblotted for the indicated proteins (TUB is tubulin). **f**, Quantification of IFNγ-R1 expression (relative to that in *EGFP*-expressing cells) by flow cytometry in D10 melanoma cells ectopically expressing either *EGFP* (control), *IFNGR1* or *JAK1*. **g**, Results of qPCR analysis for the mRNA expression of *IFNGR1* (relative to *RPL13* expression) in D10 cells expressing either *EGFP, IFNGR1* or *JAK1. IFNGR1* expression was normalized to that in *EGFP*-expressing cells. **h**, Immunoblot of either parental D10 melanoma cells, D10 *IFNGR1*-KO clones or *JAK1*-KO clones expressing either sgCtrl or sg*STUB1*. Whole cell lysates were blotted for the indicated proteins (TUB is tubulin). **i**, Quantification of IFNγ-R1 protein levels (relative to loading control and normalized to D10 parental sgCtrl-expressing cells) from (**i**). Mean±SD in (**b**), ****p<0.0001, multiple t-tests for three biological replicates, Mean±SD in (**f**): ****p<0.0001, ordinary one-way ANOVA for three biological replicates with Tukey post hoc testing. Mean±SD in (**g**): n.s. p=0.8001, ****p<0.0001, ordinary one-way ANOVA for three biological replicates with Dunnett’s post hoc testing. Mean±SD in (**i**): ***p=0.0004, ****p<0.0001, ordinary one-way ANOVA for three immunoblots with Tukey post hoc testing.

This broad effect prompted us to mechanistically dissect how STUB1 regulates IFNγ-R1 expression. qPCR analysis for *IFNGR1* showed that its transcript levels were indistinguishable between WT and *STUB1*-deficient cells (**Supplementary Fig. 2a**). Therefore, we focused our attention on a post-transcriptional mode of regulation. We first determined in which cellular compartment STUB1 regulates IFNγ-R1 expression. Cell lysates of *STUB1*-proficient and *STUB1*-deficient cells were treated with various deglycosylating enzymes. The strongest increase in IFNγ-R1 was seen in the high molecular weight, Endo-H-resistant species of IFNγ-R1. This suggests that the regulation of IFNγ-R1 by STUB1 occurs after it passes through the endoplasmic reticulum (**Supplementary Fig. 2b, c**).

IFNγ-R1 manifested as multiple protein species that were distinguishable by SDS-PAGE analysis (**Supplementary Fig. 2b**). To determine which of these forms are located at the tumor cell surface, we performed biotin labeling and immunoprecipitation of cell-surface proteins^31^. This analysis showed that solely the high molecular weight, Endo-H-resistant, species of IFNγ-R1 resides at the plasma membrane (**Supplementary Fig. 2d**). This result implies that STUB1 specifically regulates the cell surface fraction of IFNγ-R1, which is in accordance with our flow cytometry findings.

### STUB1 destabilizes IFNγ-R1 in JAK1-dependent and JAK1-independent manners

STUB1, initially identified as a co-chaperone^32^, acts as an E3 ubiquitin ligase^33,34^ that affects protein stability by mediating proteasomal degradation^34–36^. Therefore, and in accordance with our observation that STUB1 loss does not affect *IFNGR1* mRNA levels, we hypothesized that it destabilizes the IFNγ-R1 protein. To test this, we profiled the proteomes of cells expressing either a non-targeting control sgRNA (sgCtrl) or a *STUB1*-targeting sgRNA (sg*STUB1*) by mass spectrometry. This analysis not only confirmed our observation that *STUB1* inactivation increases IFNγ-R1 levels, but it also revealed a marked increase in the abundance of the JAK1 protein (**Fig. 2c**). This finding was confirmed by the same analysis in a second cell line (**Supplementary Fig. 2e**). It was also validated by immunoblotting for IFNγ-R1 and JAK1, in two melanoma cell lines (**Fig. 2d, Supplementary Fig. 2f and g**). Similar to its regulation of IFNγ-R1 expression, STUB1 also affected JAK1 protein stability, as *JAK1* transcript levels remained unchanged by *STUB1* inactivation (**Supplementary Fig. 2h**).

While it is known that the interaction of IFNγ-R1 and JAK1 is essential for the signaling functionality of the IFNγ receptor complex^37,38^, a potential role of JAK1 in stabilizing IFNγ-R1 levels, and by extension the IFNγ receptor complex, has not been reported. We first investigated whether heightened JAK1 expression would suffice to drive increased IFNγ-R1 protein stability. Ectopically expressed *JAK1* strongly increased IFNγ-R1 protein abundance (**Fig. 2e**), which translated into increased cell surface expression (**Fig. 2f**), even more so than ectopically expressed *IFNGR1* (**Fig. 2e-g and Supplementary Fig. 2i and j**). This result suggests not only that elevated JAK1 protein levels are sufficient to stabilize IFNγ-R1 protein, but also that *JAK1* expression may be crucial in dictating the amount of IFNγ-R1 present on the cell surface; unexpectedly even more so than *IFNGR1* expression itself.

To determine whether elevated JAK1 levels in *STUB1*-inactivated cells account for the rise in IFNγ-R1 abundance, we inactivated JAK1 in a *STUB1*-deficient background (**Fig. 2h and i**). This epistasis experiment revealed that *STUB1* inactivation still led to an increase in IFNγ-R1, albeit to a lesser degree than in the presence of JAK1 (**Fig. 2h and i**). These findings together indicate that *STUB1* deficiency promotes IFNγ-R1 stabilization both in JAK1-dependent and -independent fashions: STUB1 depletion increases IFNγ-R1 levels directly, but also increases JAK1 abundance, which in turn further stabilizes IFNγ-R1.

### STUB1 drives proteasomal degradation of IFNγ receptor complex through IFNγ-R1^K285^ and JAK1^K249^ residues

Since STUB1 has been shown to mediate proteasomal degradation of client proteins^35,36^, we next asked whether increased protein levels of IFNγ-R1 and JAK1 upon STUB1 inactivation were caused by reduced proteasomal degradation. We treated either wildtype or *STUB1*-deficient cells with MG132, an inhibitor of proteasomal degradation. Western blot analysis of these cell lysates showed a significant induction of IFNγ-R1 proteins in wildtype cells upon treatment with MG132 (**Fig. 3a-c**). In contrast, whereas baseline levels of IFNγ-R1 were already increased in *STUB1*-deficient cells, there was no further induction upon MG132 treatment. A similar observation was made for JAK1 (**Fig. 3a-c**). These results were recapitulated in an additional cell line (**Supplementary Fig. 3a-c**).

**Figure 3:**
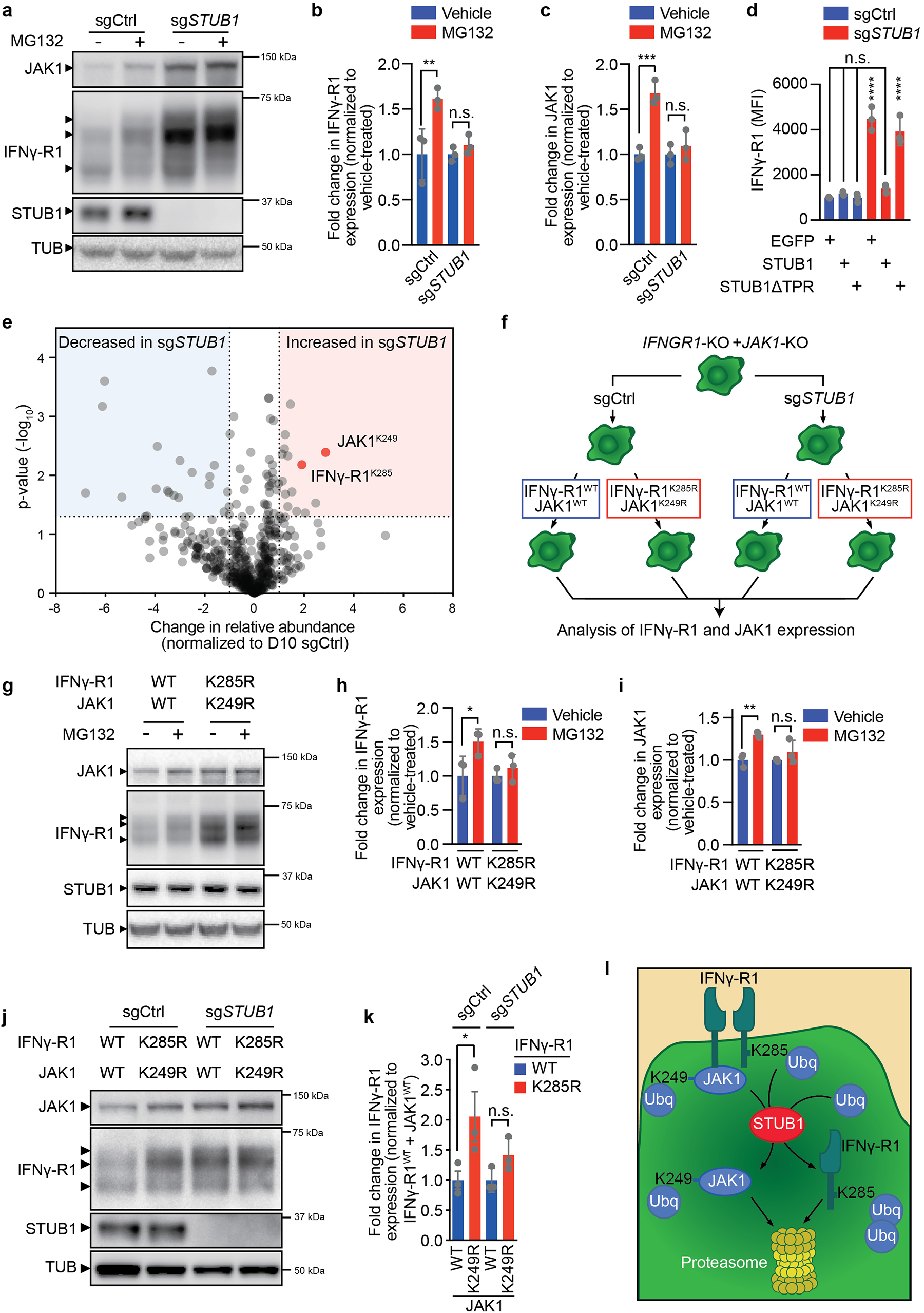
STUB1 drives proteasomal degradation of IFNγ receptor complex through IFNγ-R1^K285^ and JAK1^K249^ residues. **a**, Immunoblot of D10 melanoma cells expressing either sgCtrl or sg*STUB1* treated with either vehicle or 10 μM MG132 for four hours. Whole-cell lysates were immunoblotted for the indicated proteins (TUB is tubulin). **b**, Quantification of IFNγ-R1 protein levels (relative to loading control and normalized to vehicle-treated group) from (**a**). **c**, Quantification of JAK1 protein levels (relative to loading control and normalized to vehicle-treated group) from (**a**). **d**, MFI of IFNγ-R1 expression on D10 cells expressing sgCtrl or sg*STUB1*, which ectopically express either 3xFLAG-tagged EGFP, full length STUB1 or STUB1 lacking N-terminal residues 1-72 of the TPR domain. **e**, Relative change in K-epsilon-diglycine motif-containing peptides in sg*STUB1*-expressing cells, normalized to sgCtrl-expressing cells. Highlighted are peptides that also exhibit significant differential regulation at total protein level as assessed by global proteomic analysis (**Fig. 2c and Supplementary Figure 2e**) **f**, Schematic image depicting the reconstitution of either IFNγ-R1^WT^ and JAK1^WT^ or IFNγ-R1^K285R^ and JAK1^K249R^ ORFs in *IFNGR1*-KO + *JAK1*-KO D10 melanoma clones in either sgCtrl- or sg*STUB1*-expressing genetic background. **g**, Immunoblot of *IFNGR1*-KO + *JAK1*-KO D10 melanoma clones, reconstituted with either IFNγ-R1^WT^ and JAK1^WT^ or IFNγ-R1^K285R^ and JAK1^K249R^ ORFs. The cells were subsequently treated with 10 μM MG132 for four hours. Whole-cell lysates were immunoblotted for the indicated proteins (TUB is tubulin). **h**, Quantification of IFNγ-R1 protein levels (relative to loading control and normalized to vehicle-treated group) from (**g**). **i**, Quantification of JAK1 protein levels (relative to loading control and normalized to vehicle-treated group) from (**g**). **j**, Immunoblot on whole cell lysates of *IFNGR1*-KO + *JAK1*-KO D10 melanoma clones reconstituted with the indicated *IFNGR1* and/or *JAK1* cDNAs, as outlined in (**f**). Whole cell lysates were immunoblotted for the indicated proteins (TUB is tubulin). **k**, Fold change of IFNγ-R1 MFI (relative to *IFNGR1*-WT+*JAK1*-WT-expressing cells) in *IFNGR1*-KO + *JAK1*-KO D10 melanoma clones reconstituted with the indicated *IFNGR1* and *JAK1* cDNAs, as outlined in (**f**). Bar chart represents an excerpt from **Supplementary Fig. 3i**. **l**, Model of STUB1-mediated proteasomal degradation of IFNγ-R1 and JAK1. Mean±SD in (**b**), **p= 0.0085, n.s. p=0.8675, ordinary one-way ANOVA for three biological replicates with Tukey post hoc testing. Mean±SD in (**c**), ***p=0.0007, n.s. p=0.7936, ordinary one-way ANOVA for three biological replicates with Tukey post hoc testing. Mean±SD in (**d**), ****p<0.0001, n.s. p=0.7282, n.s. p=0.966, n.s. p=0.7154, ordinary one-way ANOVA for three biological replicates with Tukey post hoc testing. Mean±SD in (**h**), *p=0.0322, n.s. p=0.7414, ordinary one-way ANOVA for three biological replicates with Sidak post hoc testing. Mean±SD in (**i**), **p=0.0041, n.s. p=0.3570, ordinary one-way ANOVA for three biological replicates with Sidak post hoc testing. Mean±SD in (**k**), *p=0.036, n.s. p=0.9812, ordinary one-way ANOVA for three biological replicates with Tukey post hoc testing.

Proteasomal degradation by STUB1 requires its tetracorticopeptide (TPR) domain, which interacts with chaperones such as HSC70^34–36^. Therefore, we queried whether this domain is required to destabilize IFNγ-R1 and JAK1 protein levels. Reconstitution of full length STUB1 in STUB1-deficient cells resulted in reduction of IFNγ-R1 and JAK1 proteins to similar baseline levels as observed in wildtype cells (**Fig. 3d, Supplementary Fig. 3d and e**). However, STUB1-deficient cells reconstituted with a TPR-domain-deficient isoform retained elevated levels of IFNγ-R1 and JAK1 comparable to those seen in STUB1-deficient cells (**Fig. 3d, Supplementary Fig. 3d and e**). Taken together, these results indicate that STUB1 regulates protein turnover of both IFNγ-R1 and JAK1 by enabling proteasomal degradation of the latter proteins.

To understand in more detail how STUB1 mediates the proteasomal degradation of both factors, particularly which lysine residues are critical targets of STUB1, we queried the changes in the landscape of ubiquitinated proteins upon STUB1 depletion. We immunopurified peptides containing a lysine (K)-epsilon-diglycine motif; a remnant mark of ubiquitinated proteins after trypsin digestion^39^, from both wildtype and *STUB1*-inactivated cells. Then, by mass spectrometry, we identified differentially ubiquitinated lysine residues between the two genotypes. From this analysis, we learned that IFNγ-R1^K285^ and JAK1^K249^ were more frequently ubiquitinated in *STUB1*-deficient cells (**Fig. 3e**).

This raises the possibility that STUB1 specifically recognizes these ubiquitinated residues and uses them as substrates for subsequent proteasomal degradation of their respective proteins. To validate this hypothesis, we generated melanoma cell clones deficient in both *IFNGR1* and *JAK1* (*IFNGR1*-KO + *JAK1*-KO) in either a wildtype or *STUB1*-deficient background. We then reconstituted *JAK1* and *IFNGR1* either in a wildtype configuration, or in a form in which the STUB1-targeted lysine residues were mutated to arginine, thereby precluding ubiquitination events from occurring at those sites. We assessed the effects of the various mutations and genotypes on IFNγ-R1 and JAK1 protein levels by flow cytometry and Western blot (**Fig. 3f-k and Supplementary Fig. 3f-i**). This reconstitution experiment showed that preventing ubiquitination of IFNγ-R1^K285^ and JAK1^K249^ resulted in marked protein stabilization of IFNγ-R1 and JAK1 in wildtype cells (**Fig. 3g and Supplementary Fig. 3f-i, sgCtrl samples**). This increased protein stability of mutant IFNγ-R1^K285^ and JAK1^K249^ likely occurs through reduced proteasomal turnover, as MG132 treatment was unable to further stabilize IFNγ-R1 and JAK1 levels in the IFNγ-R1^K285^ and JAK1^K249^ mutants, whereas it did in wildtype cells (**Fig. 3g-i**).

To assess the reliance of STUB1 on these residues for modifying IFNγ-R1 and JAK1 stability, we continued by inactivating STUB1 in the IFNγ-R1^K285^ and JAK1^K249^ mutant cells. We analyzed IFNγ-R1 and JAK1 expression by Western blot (**Fig. 3j and Supplementary Fig. 3f-h**) and additionally assessed IFNγ-R1 expression by flow cytometry (**Fig. 3k and Supplementary Fig. 3i**). Whereas in STUB1-proficient cells, the IFNγ-R1^K285^ and JAK1^K249^ mutants resulted in increased stability of IFNγ-R1 and JAK1 (**Fig. 3j, k and Supplementary Fig. 3f-i**), they were unable to further increase IFNγ-R1 and JAK1 in a *STUB1*-KO background (**Fig. 3j, k and Supplementary Fig. 3g and h**). This finding suggests that STUB1 recognizes and requires the ubiquitinated residues IFNγ-R1^K285^ and JAK1^K249^ to target their parent proteins, IFNγ-R1 and JAK1, for proteasomal degradation (**Fig. 3l**).

### *STUB1* inactivation sensitizes melanoma cells to cytotoxic T cells through amplified IFNγ signaling

Having established that STUB1 regulates IFNγ-R1 and JAK1 expression under homeostatic conditions, we next asked whether this regulation affects receptor complex stability during active IFNγ signaling. Whereas wildtype tumor cells moderately upregulated IFNγ-R1 expression upon treatment with increasing amounts of IFNγ, *STUB1*-deficient cells further elevated IFNγ-R1 protein levels, particularly the heavier, cell-surface isoforms (**Fig. 4a**). We also observed this altered IFNγ response in *STUB1*-deficient cells with downstream mediators of IFNγ signaling, as illustrated by an accelerated and robust onset of STAT1 phosphorylation upon IFNγ treatment in STUB1-depleted cells (**Fig. 4b**). This altered signaling translated into enhanced transcription of IFNγ-responsive genes, such as *CD274* (encoding PD-L1; **Fig. 4c**) and *IDO1* (**Supplementary Fig. 4a**). We confirmed this observation at the protein level (**Supplementary Fig. 4b and c**).

**Figure 4:**
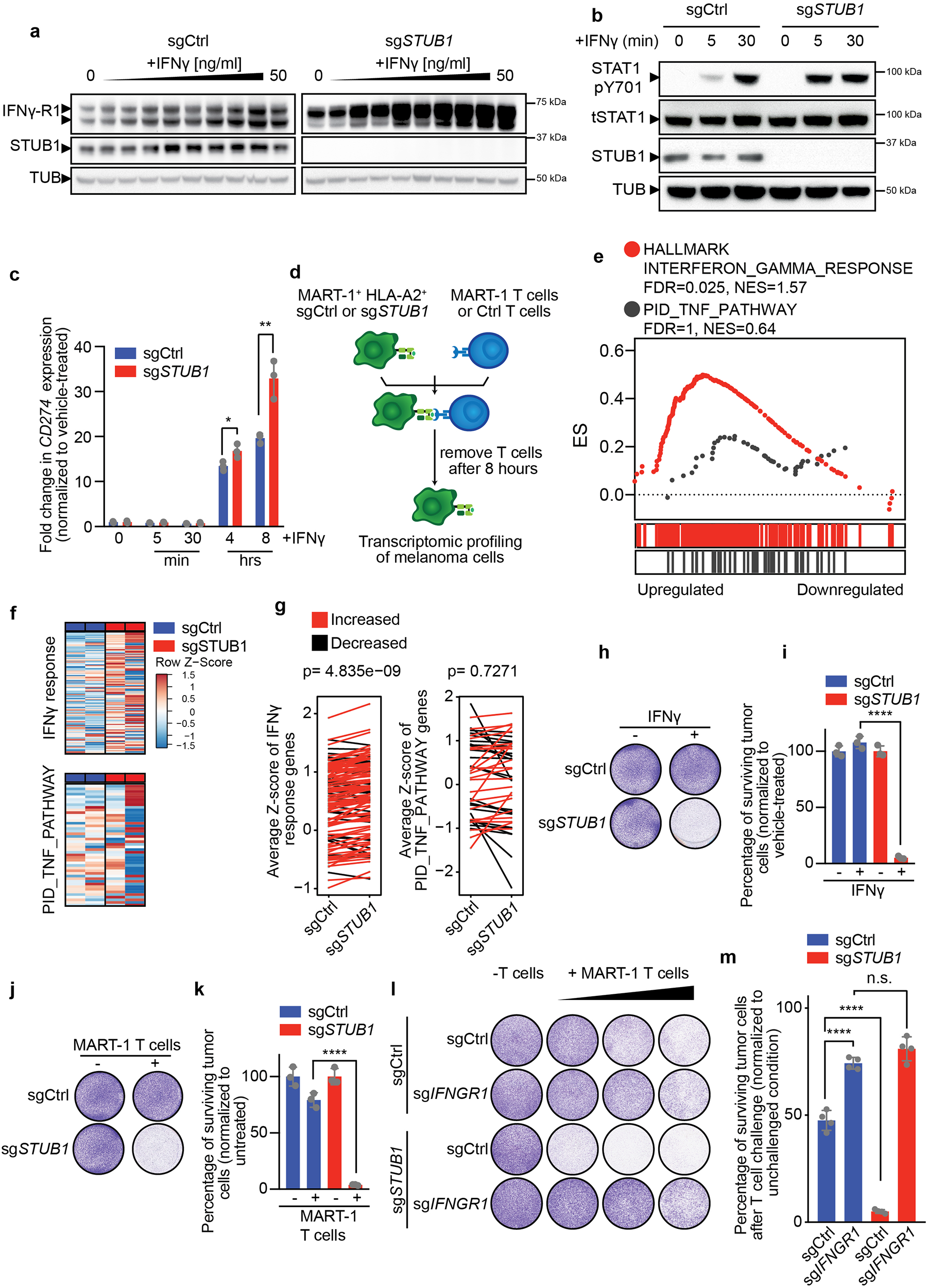
*STUB1* inactivation sensitizes melanoma cells to cytotoxic T cells through amplified IFNγ signaling. **a**, Immunoblots of D10 melanoma cells expressing sgCtrl or sg*STUB1*, treated with a two-fold serial dilution of IFNγ (starting at 50 ng/ml) for 30 minutes. Same protein amounts were loaded on two separate gels and whole cell lysates were immunoblotted for the indicated proteins (TUB is tubulin) and developed at the same time. The same exposure for the blots is shown. **b**, Immunoblot of D10 melanoma cells expressing sgCtrl or sg*STUB1*, treated with either vehicle or 50 ng/ml IFNγ for the indicated duration. Whole cell lysates were immunoblotted for phosphorylated-tyrosine-701 (pY701) of STAT1, total STAT1 (tSTAT1), STUB1 and Tubulin (TUB). **c**, qPCR analysis for the mRNA expression of *CD274* (encoding PD-L1) in D10 melanoma cells expressing either sgCtrl or sg*STUB1*, after treatment with 25 ng/ml IFNγ for the indicated duration. **d**, Schematic depiction of the experimental setup to profile the transcriptomes of D10 and SK-MEL-147 melanoma cells expressing sgCtrl or sg*STUB1*, which were co-cultured with either Ctrl or MART-1 T cells for eight hours. **e**, Gene set enrichment analysis on RNA sequencing results for D10 and SK-MEL-147 melanoma cells co-cultured with MART-1 T cells for eight hours (from **d**). **f**, Differential gene expression analysis of IFNγ response genes (derived by treating D10 and SK-MEL-147 melanoma cells with IFNγ for eight hours, depicted in **Supplementary Fig. 4d**) and PID_TNF_PATHWAY genes in D10 melanoma cells co-cultured with MART-1 T cells for eight hours. **g**, Difference in either IFNγ response gene expression or expression of PID_TNF_PATHWAY genes between sgCtrl and sg*STUB1*-expressing D10 melanoma cells following MART-1 T cell challenge for eight hours. **h**, Colony formation assay of D10 melanoma cells expressing sgCtrl or sg*STUB1* treated with either vehicle or 3 ng/ml IFNγ for five days. **i**, Quantification of colony formation assay shown in (**h**). **j**, Colony formation assay of D10 melanoma cells expressing sgCtrl or sg*STUB1* treated with either no or MART-1 T cells for 24 hours and subsequent culture for four days. **k**, Quantification of colony formation assay shown in (**j**). **l**, Colony formation assay of D10 melanoma cells expressing the indicated sgRNAs, which were co-cultured with either no or MART-1 T cells at T cell : melanoma cell ratios 1:16, 1:8 and 1:4 (left to right) for 24 hours and stained four days later. **m**, Quantification from (**l**) at a T cell : melanoma cell ratio of 1:8. Mean±SD in (**c**), **p=0.0064, *p=0.033, multiple t-tests for three biological replicates. Average Z-score of respective genes in (**g**) from two biological replicates with paired t-test. Mean±SD in (**i**), ****p<0.0001, ordinary one-way ANOVA for three biological replicates with Tukey post hoc testing. Mean±SD in (**k**), ****p<0.0001, ordinary one-way ANOVA for three biological replicates with Tukey post hoc testing. Mean±SD in (**m**), ****p<0.0001, n.s. p=0.1226, ordinary one-way ANOVA for four biological replicates with Tukey post hoc testing.

In light of these results, it was important to assess whether this hyperresponsiveness to IFNγ would also alter how STUB1-deficient tumor cells respond to T cell attack. We therefore profiled transcriptomic changes in wildtype and STUB1-deficient melanoma cells after T cell attack (**Fig. 4d**). Gene set enrichment analysis (GSEA) revealed that STUB1-depleted melanoma cells exhibit an amplified IFNγ response compared to wildtype cells (**Fig. 4e**), whereas, as a control for its specificity, genes within the TNF pathway did not show this enrichment (**Fig. 4e**). We additionally derived an experimental IFNγ response gene set from IFNγ-treated melanoma cells (**Supplementary Fig. 4d**). This gene set was significantly stronger induced in STUB1 - deficient melanoma cells challenged with cytotoxic T cells than in its control counterpart (**Fig. 4f, g**). We confirmed this effect in a second cell line (**Supplementary Fig. 4e and f**). Additionally, this effect was specific to IFNγ-signaling, as it did not occur for a TNF signaling-based gene set (**Fig. 4f, g; Supplementary Fig. 4e and f**).

Given these findings, and our previous results demonstrating that elevated IFNγ-R1 levels sensitize tumor cells to IFNγ treatment and cytotoxic T cells, we next tested whether *STUB1* inactivation induces hypersensitivity to (T cell-derived-) IFNγ. Indeed, at concentrations where wildtype melanoma cells were barely affected by IFNγ or T cell attack, *STUB1*-deficient melanoma cells were eliminated efficiently (**Fig. 4h-k and Supplementary Fig. 4g-j**). We confirmed that the sensitization to T cell attack was IFNγ-dependent, as both STUB1-deficient and wildtype cells were equally sensitive to T cell attack when lacking IFNγ-R1 expression (**Fig. 4l, m, and Supplementary Fig. 4k and l**). Collectively, these data show that the strong basal and dynamic induction of IFNγ-R1 expression by *STUB1* inactivation results in intensified IFNγ signaling and consequently, IFNγ-dependent sensitization of melanoma cells to cytotoxic T cells *in vitro*.

### STUB1 inactivation and anti-PD-1 treatment constitute a rational combination therapy approach

Having observed an enhanced sensitivity of *STUB1*-deficient melanoma cells to cytotoxic T cell pressure *in vitro* (**Fig. 4**), we next investigated whether this is recapitulated cross-species and *in vivo*. We first established *Stub1*-deficient murine melanoma cell lines in which we could reiterate our findings from human cell lines *in vitro* (**Supplementary Fig. 5a-e**). Importantly and in line with our *in vitro* data, we validated that immunogenic B16F10-dOVA tumors lacking Stub1 induced PD-L1 to a greater extent than wildtype tumors *in vivo* (**Fig. 5a-c**).

**Figure 5:**
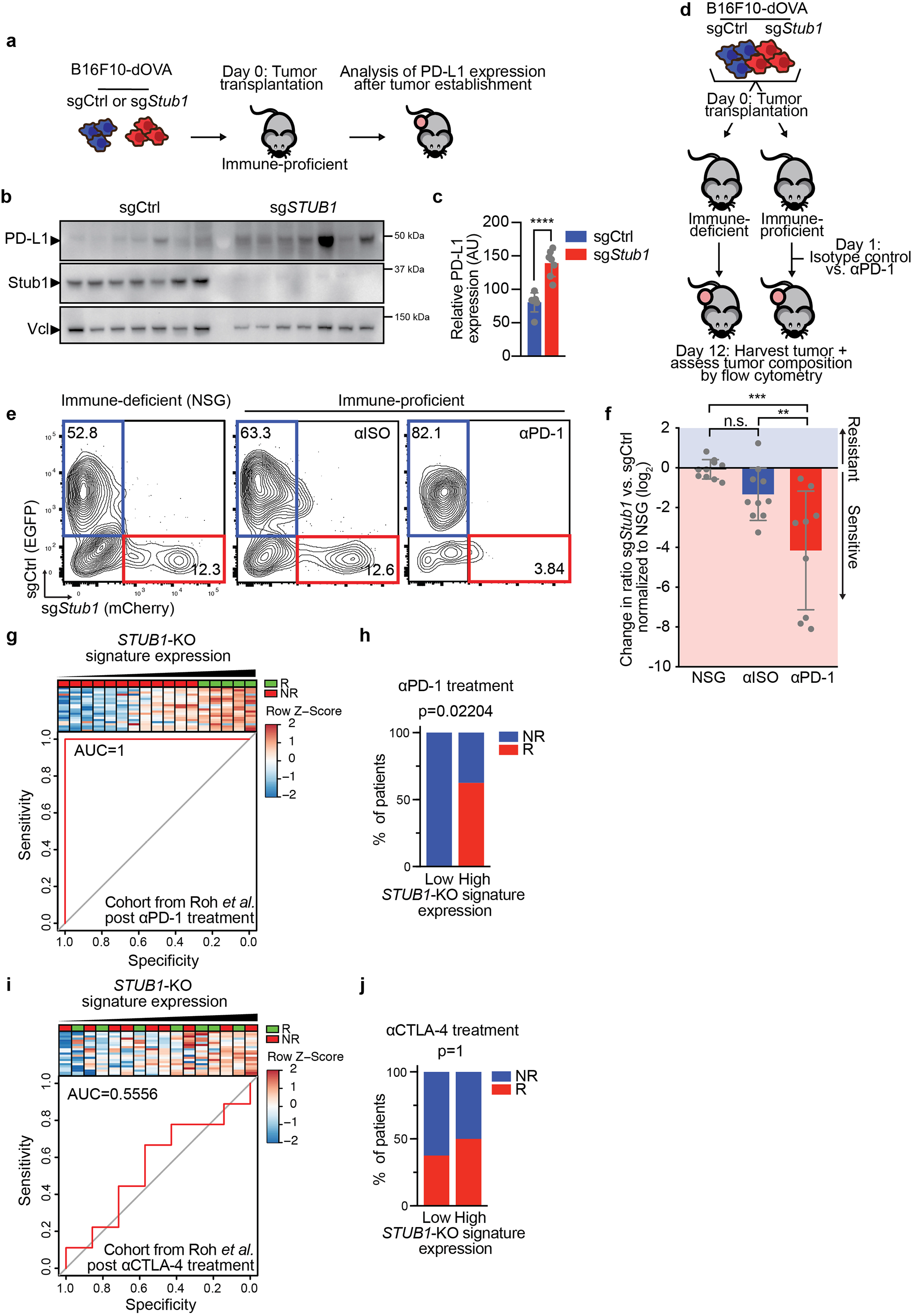
STUB1 inactivation and anti-PD-1 treatment constitute a rational combination therapy approach. **a**, Experimental outline to assess PD-L1 expression on either sgCtrl or sg*Stub1* tumors. **b**, Immunoblot of B16F10-dOVA *in vivo* tumor samples expressing either sgCtrl or sg*Stub1* (outlined in **a**) for the indicated proteins (Vcl is vinculin). **c**, Quantification of PD-L1 protein levels (relative to loading control) of tumor samples from immunoblot shown in (**b**), AU=arbitrary units. **d**, Schematic depiction of the *in vivo* competition assay modelling anti-PD-1 response with B16F10-dOVA cells expressing either sgCtrl or sg*Stub1*, which were differentially labelled with EGFP and mCherry, respectively. **e**, Flow cytometry plots from each group of the *in vivo* experiment outlined in (**d**) NSG, Isotype control-treated (αISO), anti-PD-1-treated (αPD-1). **f**, Quantification of *in vivo* competition assay outlined in (**d**). Ratios of mCherry vs. EGFP were normalized to the NSG condition. **g**, Composite plot consisting of a heat map showing melanoma patients from the post anti-PD-1 treatment cohort of Roh *et al*. 2017, sorted according to the *STUB1*-KO signature expression (average Z-score per sample), and ROC plot showing the predictive power of the *STUB1*-KO signature in this cohort. **h**, Median of *STUB1*-KO signature expression in patients from (**g**) was used to divide patients into *STUB1*-KO signature high and low-expressing groups and percentage responders and non-responders in each group was plotted. **i**, Composite plot consisting of a heat map showing melanoma patients from the post anti-CTLA-4 treatment cohort of Roh *et al*. 2017, which were sorted according to *STUB1*-KO signature expression (average Z-score per sample) and ROC plot showing the predictive power of the *STUB1*-KO signature in this cohort. **j**, Median of *STUB1*-KO signature expression in patients from (**i**) was used to divide patients into *STUB1*-KO signature high and low-expressing groups and percentage responders and non-responders in each group was plotted. Mean±SD in (**c**), **** p<0.0001, unpaired two-tailed t-test, n=7 tumors per group. Mean±SD in (**f**), *** p=0.0002, **p=0.0073, n.s. p=0.2985, ordinary one-way ANOVA with Tukey post hoc testing for n=10 in NSG and αISO and n=9 in αPD-1.

This provided a rationale to combine *Stub1* inactivation with anti-PD-1 treatment, as that combination would allow for intensified IFNγ signaling while simultaneously preventing PD-L1-mediated immune evasion. To experimentally test this, we differentially labeled wildtype and *Stub1*-deficient B16F10-dOVA cells with either EGFP or mCherry, respectively. We then mixed these cell lines in a 1:1 ratio and transplanted them into immune-deficient NSG mice, or into immune-proficient C57BL/6 mice that were subsequently treated with either an isotype control antibody or an anti-PD-1 antibody. After 12 days, tumors were harvested and the ratio between wildtype and sg*Stub1* tumor cells was assessed by flow cytometry (**Fig. 5d-f**). This analysis indicated that while there was a trend towards higher sensitivity of *Stub1*-deficient tumors to immune attack (**Fig. 5e, f**, compare NSG vs. αISO), strong depletion of *Stub1*-deficient tumors was observed only upon treatment with anti-PD-1 antibodies (**Fig. 5e, f**, compare NSG vs. αPD-1 and αISO vs. αPD-1). This finding illustrates the rationale for combining *STUB1* inhibition with anti-PD-1 therapy: it would allow for the increased susceptibility of tumors to T cell-derived IFNγ, yet at the same time block the negative effects of increased IFNγ signaling, namely increased PD-L1 levels.

To substantiate the notion that STUB1 inhibition and anti-PD-1 treatment constitute a rational treatment combination, we integrated our experimental data with clinical transcriptomic data. Based on the transcriptomic data we obtained from wildtype and *STUB1*-deficient melanoma cells after T cell attack (**Fig. 4d**), we established a *STUB1*-KO signature based on differentially upregulated genes in *STUB1*-deficient melanoma cells compared to wildtype cells upon T cell challenge (**Table 1**). We then applied this signature to transcriptomic data of melanoma patients undergoing different ICB therapies^12,40^. We found that a high *STUB1*-KO signature expression was associated with response to anti-PD-1 treatment in two cohorts (**Fig. 5g, h and Supplementary Fig. 5f, g**). Such correlation was not found for anti-CTLA-4 treatment (**Fig. 5i and j**). Importantly, these associations were not biased by the limited presence of classical IFNγ response genes in the *STUB1*-KO signature (Supplementary Fig. 5h-j). Collectively, these findings support the notion that STUB1 inactivation in combination with anti-PD-1 treatment represents a rational combinatorial treatment approach.

**Table 1:** *STUB1*-KO signature gene set. *STUB1*-KO signature gene set established through transcriptomic profiling sgCtrl or sg*STUB1*-expressing D10 and SK-MEL-147 melanoma cell lines, which were challenged with either Ctrl or MART-1-specific T cells for eight hours (outlined in Fig. 4d). The gene set is based on genes that were relatively stronger induced in sg*STUB1*-expressing cells after MART-1-specific T cell challenge.

|  |  |  |  |  |
| --- | --- | --- | --- | --- |
| *ACOD1* | *EFNA1* | *IFIH1* | *PRDX1* | *SNTB2* |
| *BIRC3* | *FST* | *IFIT2* | *PSMB8* | *SOCS3* |
| *CCL2* | *GBP1* | *IFIT3* | *RALA* | *SOD2* |
| *CD47* | *GBP2* | *IKZF2* | *RC3H1* | *THBS1* |
| *CFH* | *GBP4* | *KYNU* | *RNF145* | *TRIM22* |
| *CTSS* | *GBP5* | *LIFR* | *SAMD9L* | *XRN1* |
| *CXCL10* | *HEG1* | *OAS2* | *SAMHD1* |  |
| *CXCL11* | *HLA-B* | *OSMR* | *SELENOK* |  |
| *CXCL9* | *IDO1* | *PARP14* | *SFT2D2* |  |
| *DDX58* | *IFI44L* | *PLEKHS1* | *SLAMF8* |  |

## Discussion

Although the importance of IFNγ signaling in immunotherapy has become apparent in recent years, both experimental and preclinical studies have been largely focusing on perturbations in this pathway that contribute to tumor immunogenicity editing and immune escape^4–6,25–27,41^. Considerably less is known about the role and regulation of IFNγ-R1 cell surface expression levels, particularly whether and how increased abundance sensitizes to (T cell-derived) IFNγ. We show here that heightened IFNγ-R1 expression levels on tumor cells increases the susceptibility to T cell-derived IFNγ and its antitumor activity. This observation is underscored by clinical data strongly linking transcriptional IFNγ-dependent signaling in tumors to ICB therapy response^11,17,24^.

The relationship between IFNγ-R1 levels, IFNγ signaling and immune sensitivity raises the possibility that induction of this pathway may trigger immune responsiveness of tumor cells, something that may be therapeutically explored. Because little is known about mechanisms governing IFNγ-R1 cell surface expression, we performed an unbiased genome-wide screen and uncovered STUB1 as the most prominent hit: its loss led to increased IFNγ-receptor complex cell surface expression. STUB1 acts by mediating proteasomal degradation of its core components, IFNγ-R1 and its interaction partner JAK1. Our results suggest that STUB1 is a conserved E3 ubiquitin ligase for both IFNγ-R1 and JAK1, extending a previous observation on the ubiquitination of IFNγ-R1^42^. While STUB1 loss stabilizes cell surface IFNγ-R1, it also increases the abundance of JAK1. We show that, in turn, the increased abundance of JAK1 has a stabilizing effect on IFNγ-R1, because ectopic expression of JAK1 was sufficient to strongly stabilize IFNγ-R1. This finding was rather unexpected, given that JAK1 is believed to function solely as a kinase downstream of IFNγ-R1 following ligand engagement. Our results indicate that JAK1 in IFNγ receptor signal transduction is more influential.

Mechanistically, the identification of the critical ubiquitinated lysine residues, which STUB1 uses for ubiquitination of its IFNγ receptor targets, IFNγ-R1^K285^ and JAK1^K249^, is of relevance to understand this mode of regulation. IFNγ-R1^K285^ is located in the box 1 motif that is shared among cytokine class II receptors and is critical for JAK1 binding^43^. Conversely, JAK1^K249^ is located in the complementary FERM-domain of JAK1, enabling the binding to the box1 motif of IFNγ-R1^43^. These observations raise the possibility that JAK1 stabilizes IFNγ-R1 by masking the critical IFNγ-R1^K285^ residue prone to ubiquitination and thereby prevents subsequent STUB1-mediated proteasomal degradation. As demonstrated above, this regulatory mechanism may become even more apparent when IFNγ engages with its cognate receptor. Interestingly, this ubiquitination-mediated control of IFNγ signaling at the level of IFNγ-R1 may constitute a more common mechanism, as recently another ubiquitin ligase, FBXW7, was implicated in governing IFNγ-R1 signaling in breast cancer^44^. Our findings are complementary to this study; together they not only uncover the importance of ubiquitin-mediated IFNγ-R1 modulation but also highlight the unexpectedly broad consequences of this type of regulation, with strong effects in tumor cells ranging from heightened immune sensitivity to metastasis.

Our data suggest that as a result of IFNγ-R1 stabilization, STUB1 loss leads to enhanced IFNγ response as well as strong sensitization to cytotoxic T cell-mediated tumor cell killing. This suggests that the physiological role for STUB1 is to dampen IFNγ responses. Our findings therefore explain several previous observations. First, *STUB1* inactivation was found to sensitize tumors to immune pressure in the context of GVAX and anti-PD-1 therapy^26^; however, the underlying mechanism of this observation was unknown. Second, in a previous genome-wide loss-of-function screen for IFNγ signaling-independent tumor immune sensitizers, STUB1 was not identified as a hit^19^, highlighting its specific role as modulator of IFNγ signaling. Third, STUB1 was identified as a regulator of IFNγ-induced PD-L1 expression^45^. It was postulated that STUB1 directly mediates proteasomal degradation of PD-L1. However, we demonstrate that, instead, STUB1 acts as a modulator of IFNγ signaling and thus indirectly modulates PD-L1 expression.

In clinical trials, PD-1 blockade is now being combined with a genuine plethora of secondary treatments, although the rationale is not always fully clear from the available experimental evidence^46^. We show that STUB1 loss leads to an enhanced IFNγ-dependent transcriptional program. From a therapeutic point of view this could be beneficial, because several IFNγ target genes, such as HLA, contribute to tumor eradication. However, also PD-L1 represents an established IFNγ target, which we confirm here, and this constitutes an immune-protective tumor trait. Our observations, therefore, provide a clear rationale for combining STUB1 perturbation with PD-1 blockade. Indeed, we show that STUB1 deficiency in tumors synergizes with anti-PD-1 treatment in a murine model of melanoma. Collectively, our results therefore merit clinical exploration of inhibiting STUB1 in combination with PD-1 blockade, which will require the development of a pharmacologic inhibitor.

## Materials and Methods

### Cell lines used in the study

The human D10 (female), SK-MEL-23 (female), SK-MEL147 (female), A375 (female), SK-MEL-28 (male), BLM-M (male), 451Lu (male), A101D (male), LCLC-103H (male), HCC-4006 (male), RKO (unspecified), 8505C (female) and HEK293T (female) cell lines were obtained from the internal Peeper laboratory stock, as was the murine B16F10-OVA (male) cell line. The murine D4M.3A (male) cell line was obtained from the Blank laboratory. All cell lines were tested monthly by PCR to be negative for mycoplasma infection.

### MART-1 T cell generation

MART-1 retrovirus was made using a producer cell line as described previously^47^. Peripheral blood mononuclear cells (PBMCs) were isolated from healthy donor buffy coats (Sanquin, Amsterdam, the Netherlands) by density gradient centrifugation using Lymphoprep (Stem cell technologies, #07801). CD8 T cells were purified from the PBMC fraction using CD8 Dynabeads (Thermo Fisher Scientific, 11333D) according to manufacturer’s instructions. The isolated CD8 T cells were activated for 48 hours on non-tissue culture-treated 24-well-plates, which had been coated with anti-CD3 and anti-CD28 activating antibodies overnight (eBioscience, 16-0037-85, 16-0289-85, each 5 μg per well) at a density of 2×10^6^ cells per well. After 48 hours 2×10^6^ cells were harvested and mixed with the MART-1 virus at a 1:1 ratio and plated on a non-tissue culture-treated 24-well-plate, which had been coated with Retronectin overnight (Takara Bio, TB T100B, 25 μg per well). Spinfection was performed for two hours at 2000g. 24 hours following spinfection, MART-1 CD8 T cells were harvested and cultured for seven days, after which the transduction efficiency was assessed by flow cytometry using anti-mouse TCRβ (BD Bioscience, 553174). CD8 T cells were cultured in RPMI (Gibco, 11879020) containing 10% human serum (One Lamda, A25761), 100 units/ml penicillin, 100 μg per ml Streptomycin, 100 units/ml IL-2 (Proleukin, Novartis), 10 ng/ml IL-7 (ImmunoTools, 11340077) and 10 ng/ml IL-15 (ImmunoTools, 11340157). Following retroviral transduction, cells were maintained in RPMI containing 10% fetal bovine serum (Fisher Scientific, 15605639) and 100 units per ml IL-2.

### In vitro tumor competition assay

IFNγ-R1^Low^ and IFNγ-R1^High^-expressing tumor cells were labelled with CellTrace CFSE Cell Proliferation Dye (CFSE, Thermo Fisher Scientifc, C34554) or CellTrace Violet Cell Proliferation Dye (CTV, Thermo Fisher Scientific, C34557) according to manufacturer’s instructions. The labeled tumor cells were mixed in a 1:1 ratio and 4×10^6^ cells were seeded per 10 cm dish (Greiner). The tumor cell mix was subsequently challenged three times for 24 hours with either MART-1 T cells or control T cells at a 1:8 ratio. In parallel, the tumor cell mix was treated with either 25 ng/ml IFNγ or vehicle for five days. The surviving tumor cell fraction was analyzed for CFSE and CTV staining by flow cytometry 24 hours after the final T cell challenge or after five days of IFNγ treatment.

### IFNγ-induced PD-L1 and MHC class I expression

Tumor cells were seeded in 24-well-plates at a density of 3×10^5^ cells per well and treated either with a serial dilution series of IFNγ (PeproTech, 300-02) (starting at 50 ng/ml in two-fold dilution steps) or vehicle for 24 hours. The cells were harvested after treatment and stained for PD-L1 (eBioscience, 12-5983-42) and MHC class I (R&D Systems, FAB7098G). Induction of the respective proteins was analyzed by flow cytometry.

### Lentiviral transductions

HEK293T cells were co-transfected with pLX304 plasmids containing constructs of interest and the packaging plasmids pMD2.G (Addgene, #12259) and psPAX (Addgene, #12260) using polyethylenimine. 24 hours after transfection, the medium was replaced with OptiMEM (Thermo Fisher, 31985054) containing 2% fetal bovine serum. Another 24 hours later, lentivirus-containing supernatant was collected, filtered and stored at −80°C. Tumor cells were lentivirally transduced by seeding 5×10^5^ cells per well in a 12-well plate (Greiner), adding lentivirus at a 1:1 ratio. After 24 hours the virus-containing medium was removed and transduced tumor cells were selected with antibiotics for at least seven days.

### Sort-based genome-wide CRISRP/Cas9 knockout screen

D10 and SK-MEL-23 melanoma cells were first transduced to stably express Cas9 (lentiCas9-Blast, Addgene, #52962) and selected with blasticidin (5 μg/ml) for at least ten days. The respective cell lines were subsequently transduced with the human genome-wide CRISPR-KO (GeCKO, Addgene, #1000000048, #1000000049) sgRNA library at a 1000-fold representation and a multiplicity of infection of <0.3 to ensure one sgRNA integration per cell. The library transduction was performed in two replicates per cell line. Transduced cells were selected with puromycin (1μg/ml) for two days, after which library reference samples were harvested. Cells were cultured for an additional eight days to allow gene inactivation and establishment of the respective phenotype. Before sorting, a pre-sort bulk population was harvested. Library-transduced cells were then harvested and stained with anti-IFNγ-R1/CD119-APC antibody (Miltenyi Biotech, 130-099-921) for FACSorting. From the live cell population 10% of cells with the highest and 10% of cells with the lowest IFNγ-R1 expression were sorted. The sorted cells were washed with PBS and the cell pellet was snap frozen. Genomic DNA was isolated using the Blood and Cell culture MAXI Kit (Qiagen, 13362), according to manufacturer’s instructions. sgRNAs were amplified using a one-step barcoding PCR using NEBNext High Fidelity 2X PCR Master Mix (NEB, M0541L) and the following primers:

Forward primer: 5’-AATGATACGGCGACCACCGAGATCTACACTCTTTCCCTACACGACGCTCTTCCGATCTNNNNNNGGCTTTATATATCTTGTGGAAAGGACGAAACACC-3’
Reverse Primer: 5’-CAAGCAGAAGACGGCATACGAGATCCGACTCGGTGCCACTTTTTCAA-3’

The hexa-N nucleotide stretch contains a unique barcode to identify each sample following deep sequencing. MAGeCK (v0.5.6) was used to perform the analysis of the screen. To assess the depletion of core essential genes we compared the library reference sample to the pre-sorted bulk population. Putative regulators of IFNγ-R1 were identified by comparing the sgRNA abundance among the 10% highest and lowest IFNγ-R1-expressing populations and a signed robust rank aggregation (RRA) score was assigned to the respective genes. sgRNA targets with a false discovery rate (FDR) <0.25 were considered as putative hits.

### qPCR-based detection of transcriptomic differences

RNA from D10, SK-MEL-147 and SK-MEL-23 melanoma cells expressing either sgCtrl or sg*STUB1* was isolated using the Isolate II RNA Mini Kit (Bioline, BIO-52072) according to manufacturer’s instructions. cDNA was reverse transcribed using the Maxima First Strand cDNA synthesis kit (Fisher Scientific, 15273796) according to manufacturer’s instructions. cDNA samples were probed for the expression of *RPL13, IFNGR1, JAK1, CD274* and *IDO1* using the following primers:

*RPL13*: Forward: 5’- GAGACAGTTCTGCTGAAGAACTGAA-3’ Reverse: 5’- TCCGGACGGGCATGAC-3’
*IFNGR1*: Forward: 5’-CGGAAGTGACGTAAGGCCG-3’ Reverse: 5’-TTAGTTGGTGTAGGCACTGAGGA-3’
*JAK1*: Forward: 5’- TACCACGAGGCCGGGAC-3’ Reverse: 5’- AGAAGCGTGTGTCTCAGAAGC-3’
*CD274*: Forward: 5’- TGGCATTTGCTGAACGCATTT-3’ Reverse: 5’- AGTGCAGCCAGGTCTAATTGTT-3’
*IDO1*: Forward: 5’- AATCCACGATCATGTGAACCCA-3’ Reverse: 5’- GATAGCTGGGGGTTGCCTTT-3’

Gene Expression was quantified using the SensiFAST SYBR Hi-Rox Kit (Bioline, 92090) in combination with the StepOnePlus Real-Time PCR System (Thermo Fisher). Gene expression was normalized to *RPL13* expression using the ΔΔCt approach.

### T cell-melanoma cell co-culture

Depending on the melanoma cell line, 5×10^4^ to 1.2×10^5^ cells were seeded per well in 12-well plates in 0.5 ml DMEM containing 10% FBS. Melanoma cells were subsequently either co-cultured with the equivalent amount of control T cells or a serial dilution of MART-1 T cells in 0.5 ml DMEM containing 10% FBS (starting with a 1:1 ratio and two-fold dilution steps). After 24 hours T cells were removed by washing the plates with PBS, fresh culture medium was added and the melanoma cells were grown for four days. After the Ctrl T cell-treated well reached >80% confluence, the medium was removed and all wells were fixed with methanol and stained with crystal violet (0.1%) for 30 minutes.

B16F10-OVA cells were seeded at a density of 5×10^4^ cells per well in 0.5 ml DMEM containing 10% FBS in 12-well plates. OT-I T cells were then added in a two-fold serial dilution starting from 4:1 (T cell : melanoma cell) ratio in 0.5 ml DMEM containing 10% FBS. After 48 hours OT-I T cells were removed by washing the wells with PBS. The remaining melanoma cells were grown for an additional 48 hours, before being fixed with methanol and stained with crystal violet (0.1%). The crystal violet was removed and the plates were washed with water. After image acquisition, the crystal violet was suspended using a 10% acetic acid solution and the optical density of the resulting suspension was quantified.

### Protein expression analysis by immunoblot

Whole cell lysates were generated by removing culture medium and washing the adherent cells on the plate twice with PBS. The cells were then scraped, harvested in 1 ml PBS and pelleted by centrifugation at 1000g. After removing PBS, the cell pellet was resuspended into the appropriate amount of RIPA lysis buffer (50mM TRIS pH 8.0, 150mM NaCl, 1% Nonidet P40, 0.5% sodium deoxycholate, 0.1% SDS) supplemented with HALT Protease and Phosphatase inhibitor cocktail (Fisher Scientific, 78444). Lysis was performed on ice for 30 minutes. The samples were subsequently centrifuged at 17,000g and whole cell lysates were collected. The protein content of each lysate was quantified using Bio-Rad protein assay (Bio-Rad, 500-0006). Protein concentrations were equalized and immunoblot samples were prepared through addition of 4xLDS sample buffer (Fisher Scientific, 15484379) containing 10% β-Mercaptoethanol (final concentration 2.5%) and subsequent incubation of the samples at 95°C for five minutes. Proteins in lysates were size-separated using 4-12% Bis-Tris polyacrylamide-SDS gels (Life Technologies) and nitrocellulose membranes (GE Healthcare). Blots were blocked using 4% Milk powder in 0.2% Tween-20 in PBS. Blocked membranes were incubated with primary antibodies overnight. Immunoblots were developed using Super Signal West Dura Extended Duration Substrate (Thermo Fisher, 34075). Luminescence signal was captured by Amersham Hyperfilm high performance autoradiography film or by the Bio-Rad ChemiDoc imaging system. The following primary antibodies were used anti-IFNγ-R1 (Santa Cruz Biotechnology, sc-28363), anti-JAK1 (D1T6W, Cell Signaling Technology, 50996), anti-STUB1/CHIP (C3B6, Cell Signaling Technology, 2080), anti-Tubulin (DM1A, Sigma Aldrich, T9026), anti-STAT1 (D1K9Y, Cell Signaling Technology, 12994), anti-STAT1-Tyr701 (58D6, Cell Signaling Technology, 9167), anti-mouse PD-L1 (MIH5, Thermo Fisher Scientific, 14-5982-81).

### Quantification of protein expression of immunoblots

Protein expression on immunoblots was quantified on 8-bit gray-scale-transformed .tiff images of either scanned Amersham Hyperfilm MP (GE Healthcare, 28906838) or .tiff images obtained by the Bio-Rad ChemiDoc imaging system. Fiji ImageJ was used to select a region of interest for the respective proteins. Protein expression for each protein was normalized to the loading control of the respective sample.

### Biotin labeling of cell surface proteins

Biotin labeling of cell surface proteins was performed according to the published protocol published by Huang^31^. In brief, 2×10^6^ D10 melanoma cells were seeded in 10 cm culture dish 48 hours prior to the experiment. Cells were washed twice in ice-cold PBS/CaCl_2_/MgCl_2_ (+2.5 mM CaCl_2_, 1 mM MgCl_2_, pH 7.4). Cell surface proteins were labeled with 2 ml of 0.5 mg/ml Sulfo-NHS-SS-biotin (in PBS/CaCl_2_/MgCl_2_) on ice for 30 minutes. Labeling was quenched by washing cells three times with 3 ml of 50 mM glycine (in PBS/CaCl_2_/MgCl_2_). Cells were lysed using RIPA lysis buffer and biotinylated proteins were pulled down using Streptavidin-coated magnetic beads. Samples were size-separated using 4-12% Bis-Tris polyacrylamide-SDS gels (Life Technologies) and nitrocellulose membranes (GE Healthcare). And immunoblotted for IFNγ-R1.

### Proteome profiling

sgCtrl- and sg*STUB1*-expressing D10 and SK-MEL-147 melanoma cells (triplicates for both conditions) were lysed in 8M urea lysis buffer in the presence of cOmplete Mini protease inhibitor (Roche) and aliquots of 200 μg protein were reduced, alkylated with chloroacetamide, predigested with Lys-C (Wako) (1:75, 4h at 37°C) and trypsin digested overnight (Trypsin Gold, Mass Spectrometry Grade, Promega; 1:50 at 37°C). Peptide samples were desalted using C18 Sep-Pak cartridges (3cc, Waters) and eluted with acidic 40% and 80% acetonitrile. Dried D10 and SK-MEL-147 digests were reconstituted in 50mM HEPES buffer and replicates were labeled with 10-Plex TMT reagent (Thermo Fisher Scientific) according to the manufacturer’s instructions. Labeled samples were mixed equally for both cell lines, desalted using Sep-Pak C18 cartridges and fractionated by basic reversed-phase (HpH-RP) HPLC separation on a Phenomenex Gemini C18 analytical column (100 mm × 1 mm, particle size 3 μm, 110 Å pores) coupled to an Agilent 1260 HPLC system over a 60 minute gradient. Per cell line, fractions were concatenated to 12 fractions for proteome analysis.

Peptide fractions were analyzed by nanoLC-MS/MS on a Thermo Orbitrap Fusion hybrid mass spectrometer (Q-OT-qIT, Thermo Scientific) equipped with an EASY-NLC 1000 system (Thermo Scientific). Samples were directly loaded onto the analytical column (ReproSil-Pur 120 C18-AQ, 1.9μm, 75 μm × 500 mm, packed in-house). Solvent A was 0.1% formic acid/water and solvent B was 0.1% formic acid/80% acetonitrile. Samples were eluted from the analytical column at a constant flow of 250 nl/min in a four-hour gradient containing a 120-minute increase to 24% solvent B, a 60-minute increase to 35% B, a 40-minute increase to 45% B, 20-minute increase to 60% B and finishing with a 15-minute wash. MS settings were as follows: full MS scans (375-2000 m/z) were acquired at 120,000 resolution with an AGC target of 4×10^5^ charges and maximum injection time of 50 ms. The mass spectrometer was run in top speed mode with 3s cycles and only precursors with charge state 2-7 were sampled for MS2 using 60,000 resolution, MS2 isolation window of 1 Th, 5×10^4^ AGC target, a maximum injection time of 60 ms, a fixed first mass of 110 m/z and a normalized collision energy of 33%. Raw data files were processed with Proteome Discoverer 2.2 (Thermo Fisher Scientific) using a Sequest HT search against the Swissprot reviewed human database. Results were filtered using a 1% FDR cut-off at the protein and peptide level. TMT fragment ions were quantified using summed abundances with PSM filters requiring a S/N ≥10 and an isolation interference cutoff of 35%. Normalized protein and peptide abundances were extracted from PD2.2 and further analyzed using Perseus software (ver. 1.5.6.0)^48^. Differentially expressed proteins were determined using a t-test (cutoffs: p<0.05 and LFQ abundance difference < −0.2 ^ > 0.2).

### Ubiquitination site profiling

For ubiquitination site profiling, D10 melanoma cells expressing either a non-targeting control sgRNA (sgCtrl) or sg*STUB1* were lysed in 8M urea lysis buffer in the presence of cOmplete Mini protease inhibitor (Roche). Triplicates corresponding to 14 mg protein per sample for sgCtrl and sg*STUB1*-expressing D10 cells were reduced, alkylated with chloroacetamide, predigested with Lys-C (Wako) (1:75, 4h at 37°C) and trypsin digested overnight (Trypsin Gold, Mass Spectrometry Grade, Promega; 1:50 at 37°C). Peptide samples were desalted using C18 Sep-Pak cartridges (3cc, Waters) and eluted with acidic 40% and 80% acetonitrile. At this stage, aliquots corresponding to 200 μg protein digest were collected for proteome profiling, the remainder of the eluates being reserved for enrichment of ubiquitinated peptides. All peptide fractions were vacuum dried and stored at −80°C until further processing. Ubiquitinated peptides were enriched by immunoaffinity purification using the PTMScan Ubiquitin Remnant Motif (K-ε-GG) Kit (Cell Signaling Technology, 5562) according to the manufacturer’s instructions. Ubiquitinated peptide samples were analyzed by nanoLC-MS/MS on an Orbitrap Fusion Tribrid mass spectrometer equipped with a Proxeon nLC1000 system (Thermo Scientific) using a non-linear 210 minute gradient as described previously^49^. Raw data files were processed with MaxQuant (ver. 1.5.6.0)^50^, searching against the human reviewed Uniprot database (release 2018_01). False discovery rate was set to 1% for both protein and peptide level and GG(K) was set as additional variable modification for analysis of ubiproteome samples. Ubiquitinated peptides were quantified with label-free quantitation (LFQ) using default settings. LFQ intensities were Log2-transformed in Perseus (ver. 1.5.6.0)^48^, after which ubiquitination sites were filtered for at least two valid values (out of 3 total) in at least one condition. Missing values were replaced by an imputation-based normal distribution using a width of 0.3 and a downshift of 1.8. Differentially regulated ubiquitination sites were determined using a t-test (thresholds: p<0.05 and LFQ abundance difference < −1.0 ^ > 1.0).

### Proteasomal inhibitor treatment

Melanoma cells were seeded and grown to 80% confluence and treated with either DMSO (vehicle) or with 10 μM MG132 (Medchem Express, HY-13259) for four hours. The medium was removed four hours later, cells were washed three times with PBS and whole cell lysates were prepared as described above.

### Animal studies

All animal studies were approved by the animal ethics committee of the Netherlands Cancer Institute (NKI) and performed in accordance with ethical and procedural guidelines established by the NKI and Dutch legislation. Male mice, of either C57BL/6 (Janvier) or NSG-B2m (The Jackson Laboratory) mouse strains were used at an age of 8-12 weeks.

### In vivo tumor competition assay

B16F10-dOVA cells were lentivirally transduced with lenti-Cas9-blast to stably express Cas9 and selected with blasticidin (5 μg/ml) for at least ten days. The cells were then lentivirally transduced to stably express either sgCtrl or sg*Stub1* (lentiGuide-Puro, #52963) and cultured with puromycin (1 μg/ml) for at least ten days to allow for selection of cells with genetic inactivation of *Stub1*. Knockout efficiency was assessed by immunoblotting. sgCtrl-expressing cells were transduced to stably express EGFP (pLX304-EGFP-Blast) and sg*Stub1*-expressing cells were transduced to stably express mCherry (pLX304-mCherry-Blast). EGFP and mCherry-positive populations were sorted and cultured. Cells were mixed in a 1:1 ratio prior to injection and 5×10^5^ cell per mouse were injected into immune-deficient NSG-β2m^-/-^ (n=10, The Jackson Laboratory, 010636; RRID:ISMR_JAX:010636), or C57BL/6J mice (n=20, Janvier, C57BL/6JRj). Tumor bearing C57BL/6J mice were treated with either 100 μg/mouse isotype control antibody (Leinco Technologies, R1367) or with 100 μg/mouse anti-mouse-Pd-1 (Leinco Technologies, P372) one and six days post tumor injection. Tumors were harvested at day 12 and dissociated into single cell suspensions. Cells were subsequently stained for immune cells using anti-CD45-APC (Miltenyi, 130-102-544) and the tumor composition was analyzed by flow cytometry.

### Transcriptomic profiling of melanoma cells after T cell attack

2×10^6^ D10 and SK-MEL-147 melanoma cells were plated per dish in 10 cm cell culture dishes 48 hours prior to T cell challenge. Melanoma cells were subsequently challenged with either Ctrl or MART-1 T cells for eight hours. The T cells were removed by washing the plates with PBS. The remaining tumor cells were harvested and lysed in RLT buffer (Qiagen, 79216) and sequenced on an Illumina HiSeq2500. Fastq files were mapped to the human reference genome (Homo.sapiens.GRCh38.v77) using Tophat v2.1^51^ with default settings for single-end data. The samples were used to generate read count data using itreecount (github.com/NKI-GCF/itreecount). Normalization and statistical analysis of the expression of genes was performed using DESeq2 (V1.24.0)^52^. Centering of the normalized gene expression data was performed by subtracting the row means and scaling by dividing the columns by the standard deviation (SD) to generate a Z-score.

Differentially expressed genes between *STUB1*-deficient and wildtype cells were calculated with DESeq2^52^ using FDR<0.01. The significant genes that were up-regulated comprise the *STUB1*-KO signature (Table 1).

### External Datasets

The anti-PD-1 treated melanoma patient samples were taken from Riaz *et al*.^12^ (ENA/SRA database: PRJNA356761). Fastq files were downloaded and mapped to the human reference genome (Homo.sapiens.GRCh38.v82) using STAR(2.6.0c)^53^ in 2-pass mode with default settings for paired-end data. The samples were used to generate read count data using HTSeq-count^54^. Normalization and statistical analysis of the expression of genes was performed using DESeq2^52^. Centering of the normalized gene expression data was performed by subtracting the row means and scaling by dividing the columns by the standard deviation (SD) to generate a Z-score. Clinical data were taken from the supplementary table from the original paper. Response to ICB was based on RECIST criteria as described in the paper (Responders: CR/PR/SD, Non-Responders: PD).

Normalized gene expression data (Nanostring) and clinical data from patients treated with anti-CTLA-4 or anti-PD1 were taken from the supplementary data from Roh *et al*.^40^. Response to ICB was based on the classification from the Roh et al. manuscript (Responder or non-responder).

Heat maps were generated with matching genes between the *STUB1*-KO signature and external datasets. Samples were ordered based on the average expression of the signature (average Z-score per sample).

### GSEA

GSEAPreranked was performed using the BROAD javaGSEA standalone version (http://www.broadinstitute.org/gsea/downloads.jsp). Gene ranking was performed using the log2-fold change in gene expression between D10 and SK-MEL-147 melanoma cells expressing either sgCtrl or sg*STUB1* that were treated with MART-1 T cells for eight hours. The pre-ranked gene list was run with 1000 permutations.

## Acknowledgements

We thank all members of the Peeper and Blank laboratories as well as of the Division of Molecular Oncology and Immunology for constructive feedback and valuable input. We thank R. Mezzadra, C. Sun, T. Schumacher as well as J. Staring and T. Brummelkamp for sharing reagents and cell lines. Furthermore, we thank the flow cytometry, proteomics and sequencing core facilities as well as the animal housing facility of The Netherlands Cancer Institute for their support.

## Author contributions

G.A., D.W.V. and D.S.P. conceptualized the project, G.A. and D.W.V. performed the experiments and contributed equally to this work. O.B.B. and M.A. performed the proteomic profiling experiments. O.K. performed bioinformatic analyses for transcriptomic profiling. M.A.L., B.B. and J.B. carried out mouse experiments. D.D.A. performed experiments. J.D.L. provided wildtype and mutant *IFNGR1*-ORF constructs. M.A. and O.B.B. acknowledge support of the X-omics Initiative, part of the NWO National Roadmap for Large-Scale Research Infrastructures. G.A., D.W.V. and D.S.P. wrote the manuscript. D.S.P. supervised this study.

## Supplementary Figure Legends

**Supplementary Figure 1:**
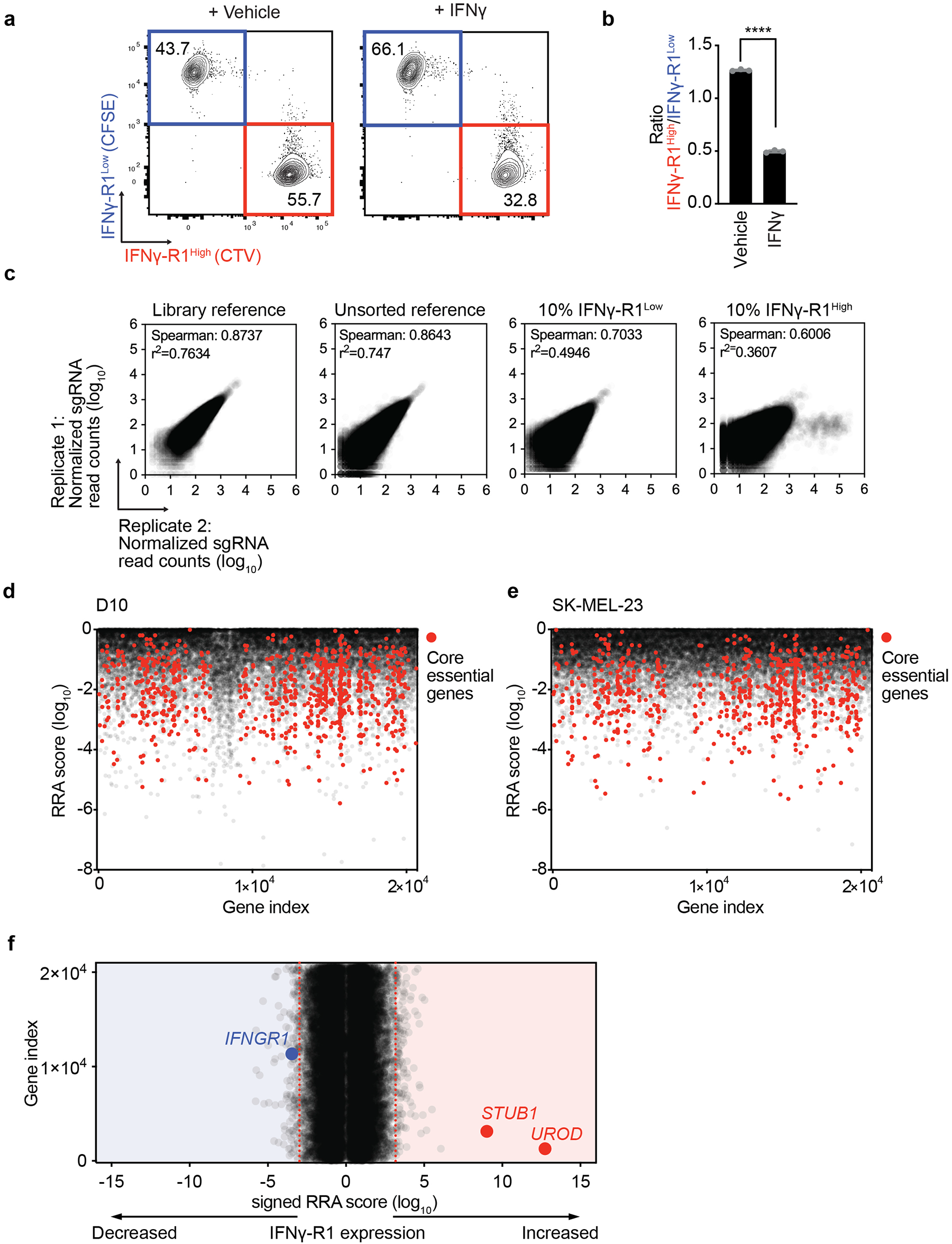
Genome-wide CRISPR/Cas9 knockout screen identifies negative regulators of IFNγ-R1 expression to modulate its cell surface abundance. **a**, Flow cytometry plot of *in vitro* competition assay of IFNγ-R1^High^ vs. IFNγ-R1^Low^ cells treated with either vehicle or 25 ng/ml IFNγ for five days. **b**, Quantification of the ratio IFNγ-R1^High^ : IFNγ-R1^Low^ in competition assay of (**a**). **c**, Correlation plots of log_10_-transformed normalized read counts of sgRNAs in genome-wide CRISPR-KO screen in D10 melanoma cell line between replicates. **d**, **e**, Log_10_-transformed RRA scores of depleted genes comparing library reference sample to unsorted bulk population in D10 (**d**) and SK-MEL-23 cells (**e**). Highlighted in red: core essential genes. y-axis: RRA score, x-axis: gene index. **f**, Results of screen outlined in (**Figure 1f**) for SK-MEL-23 cells. x-axis: signed log_10_-transformed signed MAGeCK robust rank aggregation (RRA) score for each gene; y-axis: gene index. Red dotted lines indicate FDR cutoff <0.25 for genes enriched in 10% of cells with the highest (right) or lowest (left) IFNGR1 expression. Mean±SD in (**b**), ****p<0.0001, unpaired t-test for three biological replicates.

**Supplementary Figure 2:**
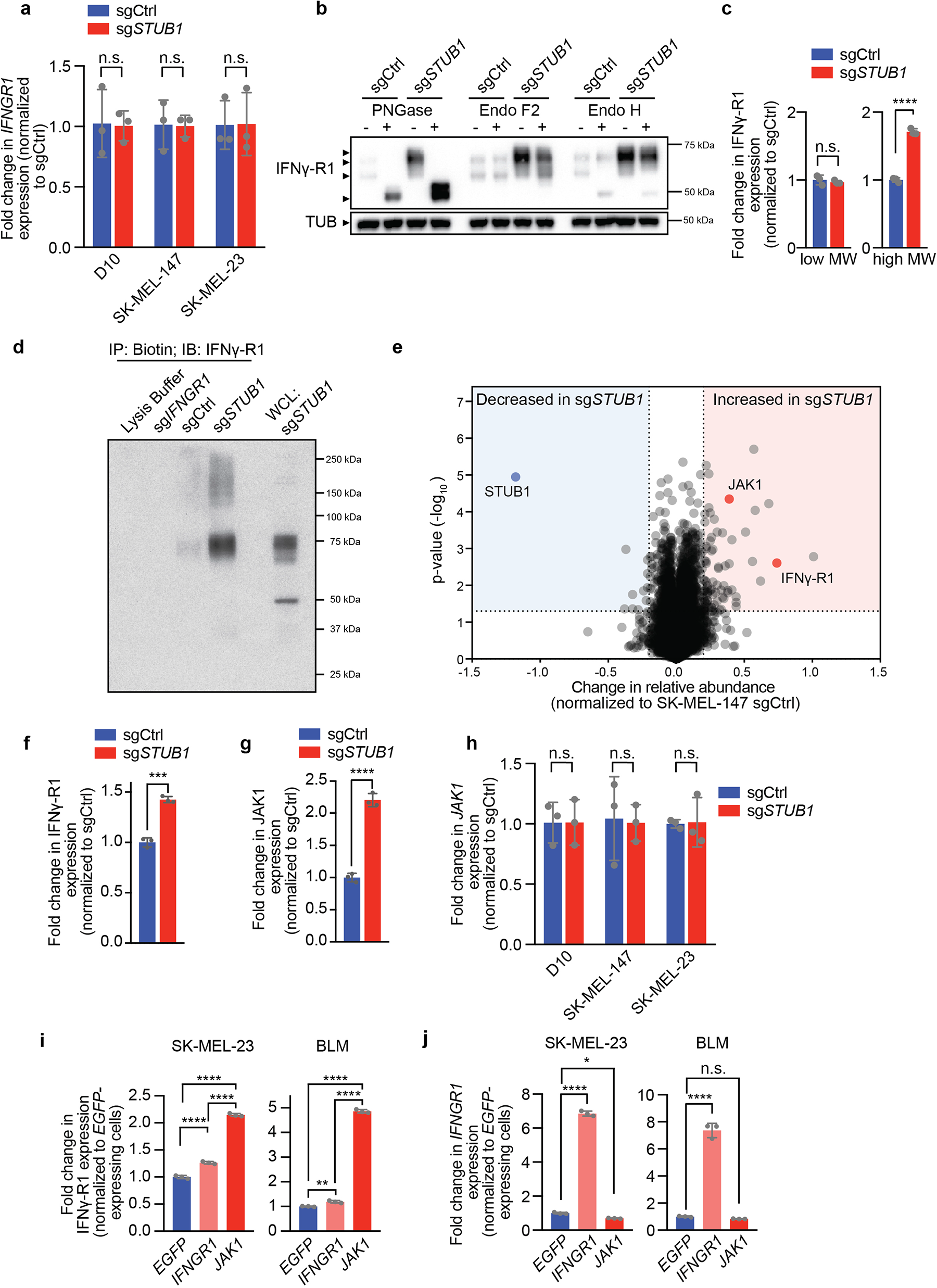
STUB1 destabilizes cell surface IFNγ-R1 in JAK1-dependent and JAK1-independent manners. **a**, Results of qPCR analysis for *IFNGR1* mRNA expression (relative to *RPL13* expression) in D10, SK-MEL-147 and SK-MEL-23 cells expressing either sgCtrl or sg*STUB1*. **b**, Immunoblot of whole cell lysates treated with the indicated deglycosylating enzymes. Whole cell lysates were collected from D10 melanoma cells expressing either sgCtrl or sg*STUB1*. Whole cell lysates were immunoblotted for IFNγ-R1 and Tubulin. **c**, Quantification of low and high molecular weight IFNγ-R1 protein levels (relative to loading control) in D10 melanoma cells from immunoblot shown in (**Figure 2d**). **d**, Immunoblot of immuno-precipitated cell surface proteins using biotin labelling in D10 melanoma clone deficient in *IFNGR1*, or D10 melanoma cell pool expressing either sgCtrl or sg*STUB1*. Following immunoprecipitation of biotin-labelled proteins, samples were immunoblotted for IFNγ-R1. The right-most lane represents 10% of the whole cell lysate of sg*STUB1*-expressing cells. **e**, Results of proteomic profiling of SK-MEL-147 melanoma cells expressing either sgCtrl or sg*STUB1*. Highlighted are the top differentially regulated proteins shared between sgCtrl and sg*STUB1*-expressing D10 and SK-MEL-147 cells (**Fig. 2c**). **f**, Quantification of IFNγ-R1 protein levels (relative to loading control) in D10 melanoma cells from immunoblot shown in (**Figure 2d**). **g**, Quantification of JAK1 protein levels (relative to loading control) in D10 melanoma cells from immunoblot shown in (**Figure 2d**). **h**, Results of qPCR analysis for *JAK1* mRNA expression (relative to *RPL13* expression) in D10, SK-MEL-147 and SK-MEL-23 cells expressing either sgCtrl or sg*STUB1*. **i**, Quantification of IFNγ-R1 expression (relative to *EGFP*-ORF-expressing cells) by flow cytometry in SK-MEL-23 and BLM-M melanoma cells expressing *EGFP*-ORF, *IFNGR1*-ORF and *JAK1*-ORF. **j**, Results of qPCR analysis for *IFNGR1* mRNA expression (relative to *RPL13* expression) in SK-MEL-23 and BLM-M melanoma cells expressing *EGFP*-ORF, *IFNGR1*-ORF and *JAK1*-ORF. Relative *IFNGR1* expression was normalized to *EGFP*-ORF-expressing cells. Mean±SD in (**a**), D10: n.s. p=0.918, SK-MEL-147: n.s. p=0.933, SK-MEL-23: n.s. p=0.968, unpaired t-tests were performed for each cell line, each three biological replicates. Mean±SD in (**c**), ****p<0.0001, n.s. p=0.5029, unpaired t-test for three biological replicates. Mean±SD in (**f**), ***p=0.0002, unpaired t-test for three biological replicates. Mean±SD in (**g**), ****p<0.0001, unpaired t-test for three biological replicates. Mean±SD in (**h**), D10: n.s. p=0.99, SK-MEL-147: n.s. p=0.877, SK-MEL-23: n.s. p=0.921, multiple t-test for three biological replicates. Mean±SD in (**i**), **p=0.0093, ****p<0.0001, ordinary one-way ANOVA for three biological replicates with Tukey post hoc testing. Mean±SD in (**j**), *p=0.0103, ****p<0.0001, n.s. p=0.7409, ordinary one-way ANOVA for three biological replicates with Dunnett post hoc testing.

**Supplementary Figure 3:**
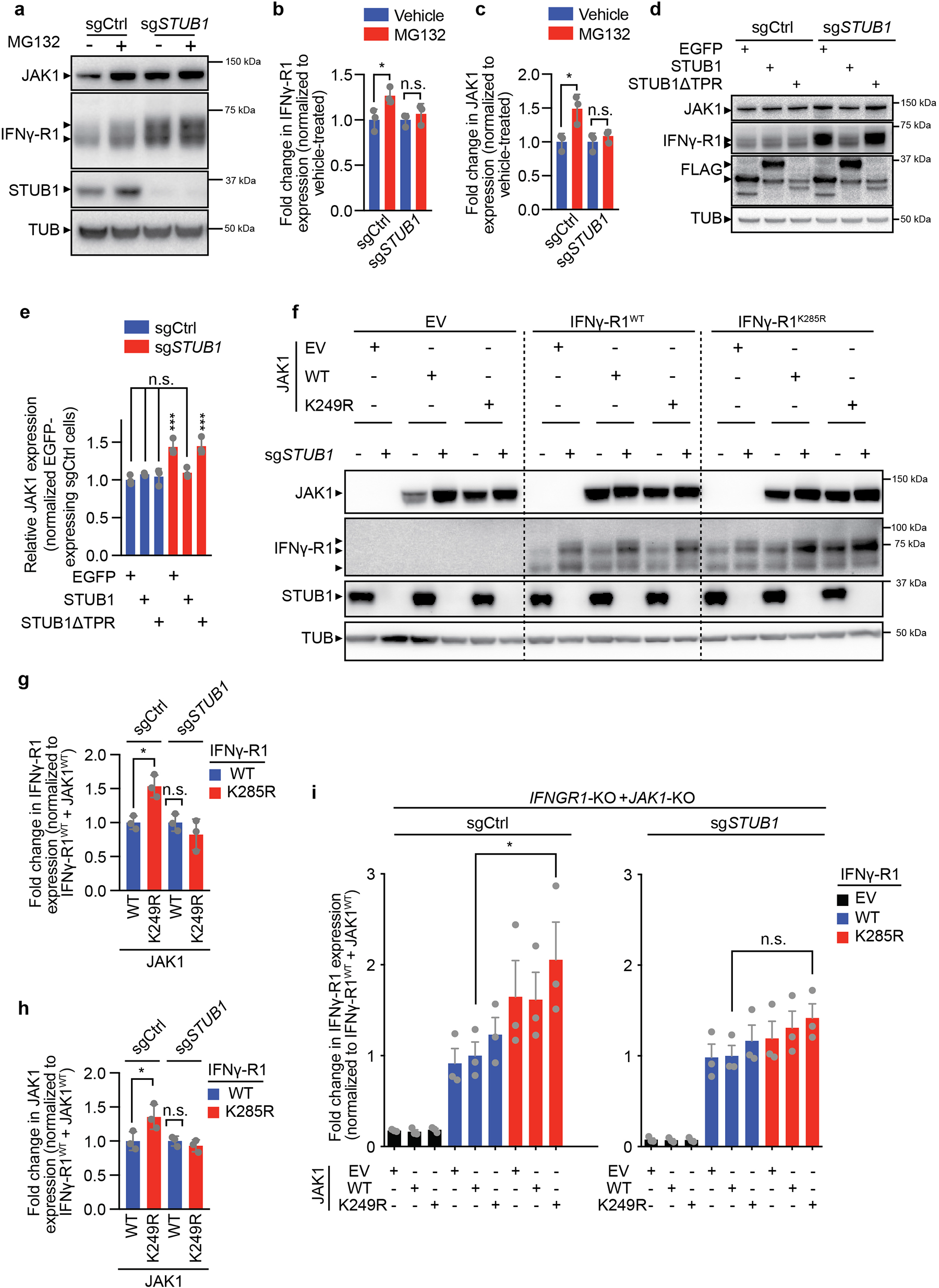
STUB1 drives proteasomal degradation of IFNγ receptor complex through IFNγ-R1^K285^ and JAK1^K249^ residues. **a**, Immunoblot of SK-MEL-147 melanoma cells expressing either sgCtrl or sg*STUB1* treated with either vehicle or 10 μM MG132 for four hours. Whole-cell lysates were immunoblotted for the indicated proteins (TUB is Tubulin). **b**, Quantification of IFNγ-R1 protein levels (relative to loading control and normalized to vehicle-treated group) from (**a**). **c**, Quantification of JAK1 protein levels (relative to loading control and normalized to vehicle-treated group) from (**a**). **d**, Immunoblot of D10 melanoma cells expressing either sgCtrl or sg*STUB1*, that ectopically express either 3xFLAG-tagged EGFP, full length STUB1 or STUB1 lacking N-terminal residues 1-72 of the TPR domain. Whole cell lysates were blotted for the indicated proteins (TUB is Tubulin). **e**, Quantification of JAK1 protein levels (relative to loading control and normalized to EGFP- and sgCtrl-expressing cells) from immunoblot depicted in (**d**). **f**, Immunoblot of whole cell lysates from *IFNGR1*-KO + *JAK1*-KO D10 melanoma clones reconstituted with the indicated *IFNGR1* and *JAK1* cDNAs, for the indicated proteins (TUB is Tubulin). **g**, Quantification of IFNγ-R1 protein levels on immunoblot in **Figure 3j** (relative to loading control and normalized to IFNγ-R1^WT^ and JAK1^WT^-expressing cells) in *IFNGR1*-KO + *JAK1*-KO D10 melanoma clones expressing either IFNγ-R1^WT^ and JAK1^WT^ or with IFNγ-R1^K285R^ and JAK1^K249R^. **h**, Quantification of JAK1 protein levels on immunoblot in **Figure 3j** (relative to loading control and normalized to IFNγ-R1^WT^ and JAK1^WT^-expressing cells) in *IFNGR1*-KO + *JAK1*-KO D10 melanoma clones expressing either IFNγ-R1^WT^ and JAK1^WT^ or with IFNγ-R1^K285R^ and JAK1^K249R^. **i**, Quantification of IFNγ-R1 expression by flow cytometry in *IFNGR1*-KO + *JAK1*-KO D10 melanoma clones reconstituted with the indicated *IFNGR1* and *JAK1* cDNAs (outlined in **Figure 3f**), shown as fold-change of IFNγ-R1 MFI relative to IFNγ-R1^WT^ + JAK1^WT^-expressing cells for each respective genotype. EV = empty vector control. Mean±SD in (**b**), *p=0.0435, n.s. p=0.8357, ordinary one-way ANOVA for three biological replicates with Tukey post hoc testing. Mean±SD in (**c**), *p=0.0138, n.s. p=0.8846, ordinary one-way ANOVA for three biological replicates with Tukey post hoc testing. Mean±SD in (**e**), ***p=0.004, ***p=0.003, n.s. p=0.7405, p=9996, p=0.972, ordinary one-way ANOVA for three biological replicates with Tukey post hoc testing. Mean±SD in (**g**), *p=0.0156, n.s. p=0.5704, ordinary one-way ANOVA for three immunoblots with Tukey post hoc testing. Mean±SD in (**h**), *p=0.0366, n.s. p=0.9068, ordinary one-way ANOVA for three biological replicates with Tukey post hoc testing. Mean±SD in (**i**), *p=0.036, n.s. p=0.9812, ordinary one-way ANOVA for three biological replicates, with Tukey post hoc testing.

**Supplementary Figure 4:**
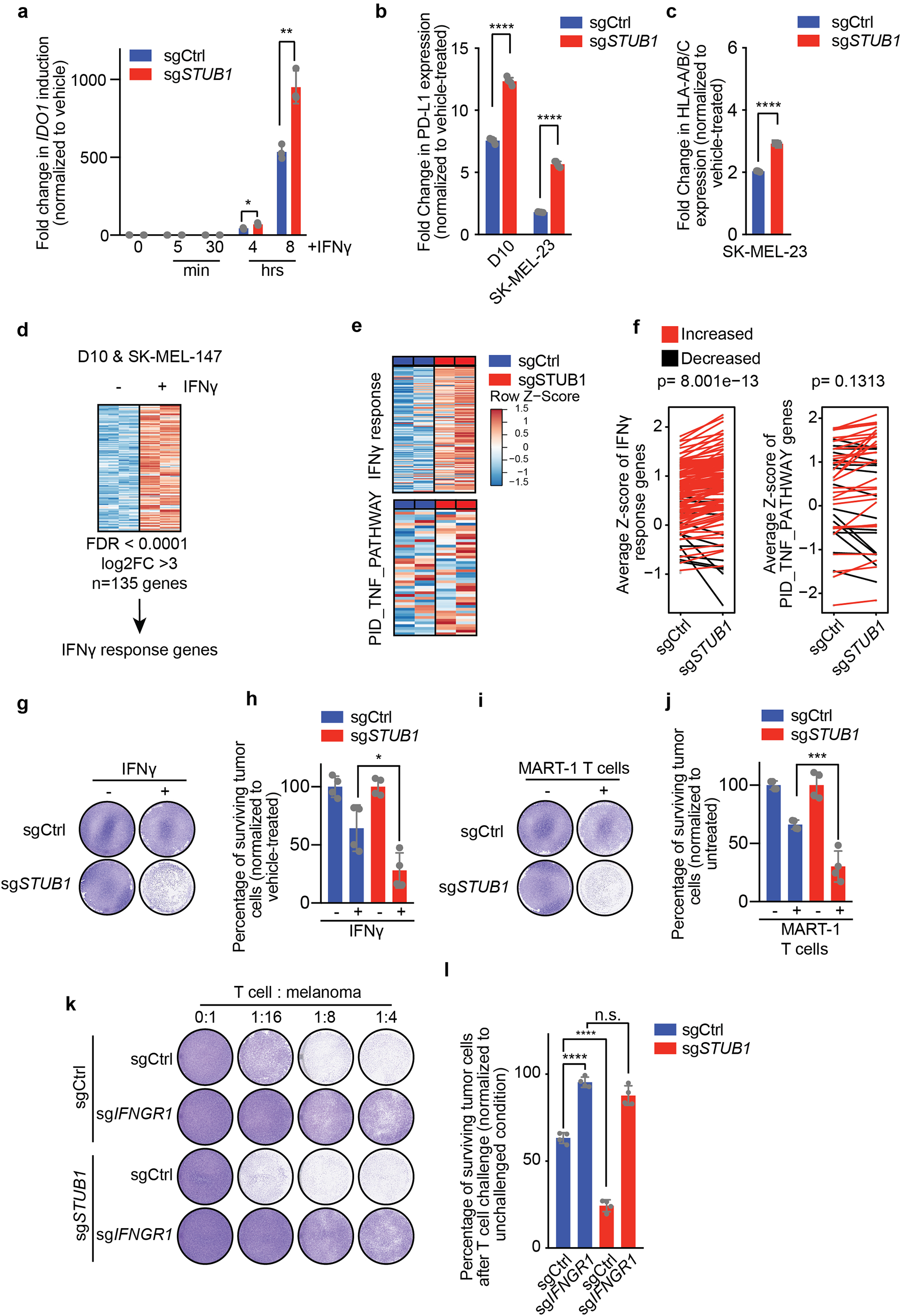
*STUB1* inactivation sensitizes melanoma cells to cytotoxic T cells through amplified IFNγ signaling. **a**, qPCR analysis for *IDO1* mRNA expression in D10 melanoma cells expressing either sgCtrl or sg*STUB1*, which were treated with 25 ng/ml IFNγ for the indicated duration. **b**, Flow cytometry analysis of IFNγ-induced PD-L1 expression on cells expressing either sgCtrl or sg*STUB1* after 24 hours treatment with 5 ng/ml IFNγ for D10 cells and 0.5 ng/ml IFNγ for SK-MEL-23 cells. **c**, Flow cytometry analysis of IFNγ-induced HLA-A/B/C expression on SK-MEL-23 melanoma cells expressing either sgCtrl or sg*STUB1* after 24 hours treatment with 0.5 ng/ml IFNγ for SK-MEL-23. **d**, Differential gene expression of D10 and SK-MEL-147 melanoma cells lines after treatment with IFNγ for eight hours was used to derive an IFNγ response gene set. **e**, Differential gene expression analysis of IFNγ response genes (derived by treating D10 and SK-MEL-147 melanoma cells with IFNγ for eight hours, depicted in **d**) and PID_TNF_PATHWAY genes in SK-MEL-147 melanoma cells co-cultured with MART-1 T cells for eight hours. **f**, Difference in either IFNγ response gene expression or expression of PID_TNF_PATHWAY genes between sgCtrl and sg*STUB1*-expressing SK-MEL-147 melanoma cells following MART-1 T cell challenge for eight hours. **g**, Colony formation assay of SK-MEL-147 melanoma cells expressing sgCtrl or sg*STUB1* treated with either vehicle or 50 ng/ml IFNγ for five days. **h**, Quantification of colony formation assay shown in (**g**). **i**, Colony formation assay of SK-MEL-147 melanoma cells expressing sgCtrl or sg*STUB1* treated with either no or MART-1 T cells for 24 hours and subsequent culture for four days. **j**, Quantification of colony formation assay shown in (**i**). **k**, Colony formation assay of SK-MEL-147 melanoma cells expressing the indicated sgRNAs that were co-cultured with either no T cell or MART-1 T cells at T cell : melanoma cell ratios 1:16, 1:8 and 1:4 (left to right) for 24 hours and subsequent culture for four days. **l**, Quantification of crystal violet stained colony formation assays from (**k**) at a T cell : melanoma cell ratio of 1:16. Mean±SD in (**a**), **p=0.0034, *p=0.012, multiple t-tests for three biological replicates. Mean±SD in (**b**), ****p<0.0001 for SK-MEL-23, ****p<0.0001, unpaired t-test for five biological replicates. Mean±SD in (**c**), ****p<0.0001, unpaired t-test for five biological replicates. Average Z-score of respective genes in (**g**) from two biological replicates with paired t-test. Mean±SD in (**h**), *p=0.0132, ordinary one-way ANOVA for four biological replicates with Tukey post hoc testing. Mean±SD in (**j**), ***p=0.0006, ordinary one-way ANOVA for four biological replicates with Tukey post hoc testing. Mean±SD in (**l**), n.s. p=0.0713, ****p<0.0001, ordinary one-way ANOVA for four biological replicates with Tukey post hoc testing.

**Supplementary Figure 5:**
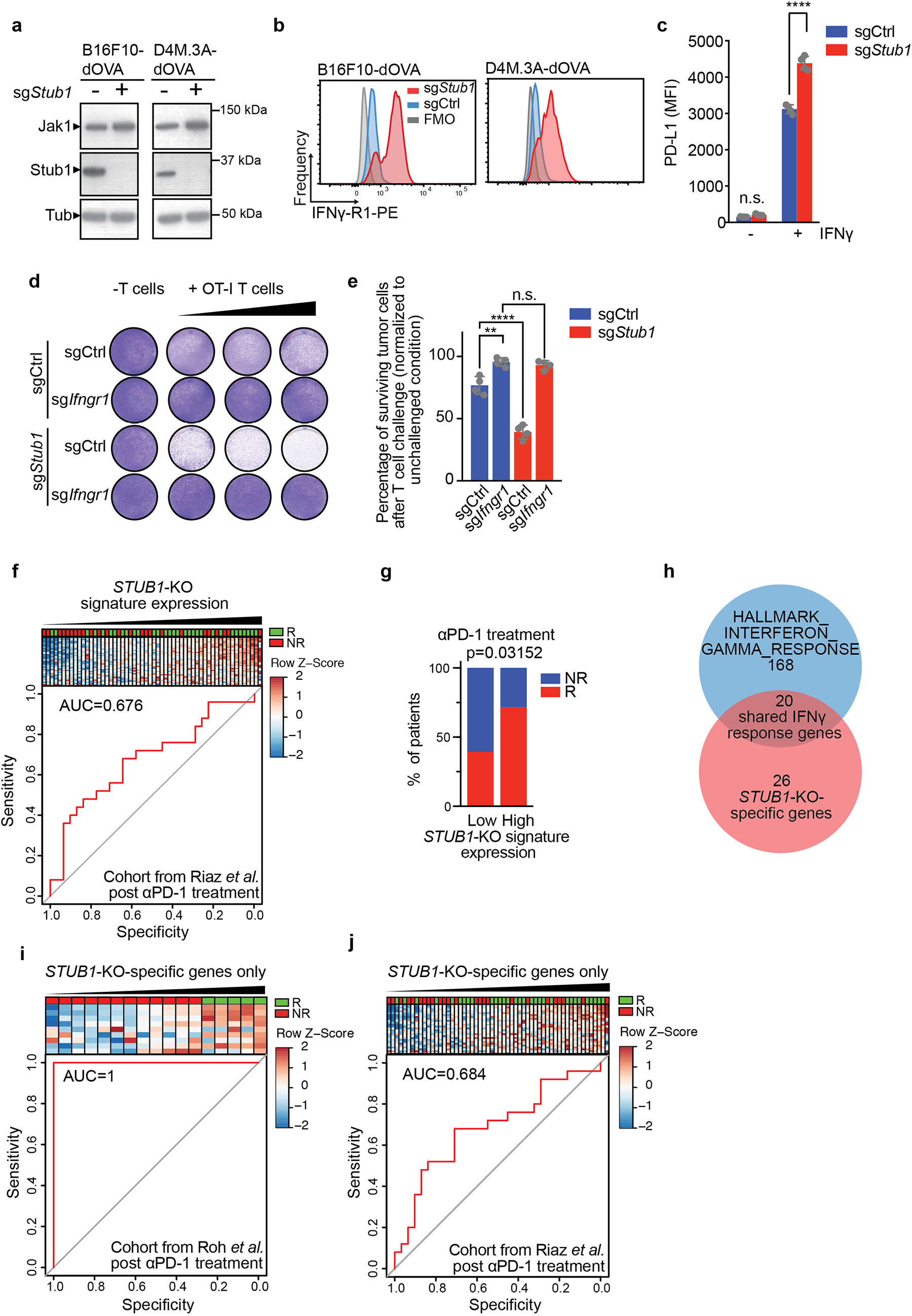
STUB1 inactivation and anti-PD-1 treatment constitute a rational combination therapy approach. **a**, Immunoblot of murine melanoma cell lines expressing either sgCtrl or sg*Stub1*. Whole cell lysates were blotted for the indicated proteins (TUB is Tubulin). **b**, Flow cytometry histograms showing Ifngr1 expression in indicated murine melanoma cell lines expressing either sgCtrl (blue) or sg*Stub1* (red). FMO (grey) = Fluorescence minus one, PE=Phycoerythrin. **c**, Flow cytometry analysis of IFNγ-induced PD-L1 expression in B16F10-dOVA cells expressing either sgCtrl or sg*Stub1*. Cells were treated with 12 ng/ml murine IFNγ for 24 hours. **d**, Colony formation assay of B16F10-dOVA melanoma cells expressing the indicated sgRNAs and co-cultured with either no T cells or OT-I T cells at T cell : melanoma cell ratios 1:1, 2:1 and 4:1 (left to right). **e**, Quantification from (**d**) at a T cell : melanoma cell ratio of 4:1. **f**, Same analysis as in **Figure 5g**, for melanoma patients from the post αPD-1-treatment cohort of Riaz *et al*. 2017. **g**, Same analysis as in **Figure 5h**, for the post αPD-1-treatment cohort of Riaz *et al*. 2017 **h**, Venn diagram depicting the overlap between the HALLMARK_INTERFERON_GAMMA_RESPONSE gene set and the *STUB1*-KO signature gene set. **i**, Same analysis as in **Figure 5g**, for the post αPD-1-treatment cohort of Roh *et al*. 2017 using the expression of the 26 genes specific to the *STUB1*-KO signature (outlined in **h**). **j**, Same analysis as in **Figure 5g**, for the post αPD-1-treatment cohort of Riaz *et al*. 2017 using the expression of the 26 genes specific to the *STUB1*-KO signature (outlined in **h**). Mean±SD in (**c**), ****p p<0.0001, n.s. p=0.8893, ordinary one-way ANOVA for four biological replicates with Tukey post hoc testing. Mean±SD in (**e**), **p=0.0012, ****p<0.0001, n.s. p=0.9012, ordinary one-way ANOVA for four biological replicates with Tukey post hoc testing.

